# Altered visual cortex excitatory/inhibitory ratio following transient congenital visual deprivation in humans

**DOI:** 10.1101/2024.04.18.590147

**Authors:** Rashi Pant, Kabilan Pitchaimuthu, José Ossandón, Idris Shareef, Sunitha Lingareddy, Jürgen Finsterbusch, Ramesh Kekunnaya, Brigitte Röder

## Abstract

Non-human animal models have indicated that the ratio of excitation to inhibition (E/I) in neural circuits is experience dependent and changes across development. Here, we assessed 3T Magnetic Resonance Spectroscopy (MRS) and electroencephalography (EEG) markers of cortical E/I ratio in ten individuals who had been treated for dense bilateral congenital cataracts, after an average of 12 years of blindness, to test for dependence of the E/I ratio in humans on early visual experience. First, participants underwent MRS scanning at rest with their eyes opened and eyes closed, to obtain visual cortex Gamma-Aminobutyric Acid (GABA+) concentration, Glutamate/Glutamine (Glx) concentration and the concentration ratio of Glx/GABA+, as measures of inhibition, excitation, and E/I ratio respectively. Subsequently, EEG was recorded to assess aperiodic activity (1-20 Hz) as a neurophysiological measure of the cortical E/I ratio, during rest with eyes open and eyes closed, and during flickering stimulation. Across conditions, congenital cataract-reversal individuals demonstrated a significantly lower visual cortex Glx/GABA+ ratio, and a higher intercept and steeper aperiodic slope at occipital electrodes, compared to age-matched sighted controls. In the congenital cataract-reversal group, a lower Glx/GABA+ ratio was associated with better visual acuity, and Glx concentration correlated positively with the aperiodic intercept in the conditions with visual input. We speculate that these findings result from an increased E/I ratio of the visual cortex as a consequence of congenital blindness, which might require commensurately increased inhibition in order to balance the additional excitation from restored visual input. The lower E/I ratio in congenital cataract-reversal individuals would thus be a consequence of homeostatic plasticity.

## INTRODUCTION

Sensitive periods are epochs during the lifespan within which effects of experience on the brain are particularly strong (Knudsen, 2004). Non-human animal work has established that structural remodelling (Bourgeois, 1997) and the development of local inhibitory neural circuits strongly link to the timing of sensitive periods (Gianfranceschi et al., 2003; Hensch et al., 1998; Hensch & Bilimoria, 2012; Hensch & Fagiolini, 2004; Takesian & Hensch, 2013). Early visual experience has been shown to fine-tune local inhibitory circuits (Benevento et al., 1992; Chattopadhyaya et al., 2004; Gandhi et al., 2008; Toyoizumi et al., 2013), which dynamically control feedforward excitation (Tao & Poo, 2005; Wu et al., 2022). The end of the sensitive period has been proposed to coincide with the maturation of inhibitory neural circuits (Hensch, 2005; Wong-Riley, 2021; H. Zhang et al., 2018). Within this framework, neural circuit stability following sensitive periods is maintained via a balance between excitatory and inhibitory transmission across multiple spatiotemporal scales (Froemke, 2015; Haider et al., 2006; Maffei et al., 2004; Takesian & Hensch, 2013; Wu et al., 2022). Such an excitatory/inhibitory (E/I) ratio has been studied at different organizational levels, including the synaptic and neuronal levels, as well as for neural circuits (Van Vreeswijk & Sompolinsky, 1996; Wu et al., 2022).

The experience-dependence of local inhibitory circuit tuning in early development is supported by a large body of work in non-human animals. In particular, studies of the mouse visual cortex has demonstrated a disrupted tuning of local inhibitory circuits as a consequence of lacking visual experience at birth (Hensch & Fagiolini, 2004; Levelt & Hübener, 2012). In addition, dark-reared mice have been shown to have increased spontaneous neural firing in adulthood (Benevento et al., 1992) and a reduced magnitude of inhibition, particularly in layers II/III of the visual cortex (Morales et al., 2002), suggesting an overall higher level of excitation.

Human neuroimaging studies have similarly demonstrated that visual experience during the first weeks and months of life is crucial for the development of visual neural circuits (Baroncelli et al., 2011; Lewis & Maurer, 2005; Maurer & Hensch, 2012; Röder et al., 2021; Röder & Kekunnaya, 2021; Singh et al., 2018). As studies manipulating visual experience are impossible in human research, much of our understanding of the experience-dependence of visual circuit development comes from patients who underwent a transient period of congenital blindness due to dense bilateral congenital cataracts. If human infants born with dense bilateral cataracts are treated later than a few weeks from birth, they suffer from a permanent reduction of visual acuity (Birch et al., 1998; Khanna et al., 2013), stereovision (Birch et al., 1993; Tytla et al., 1993), and impairments in higher-level visual functions such as face perception (Le Grand et al., 2001; Putzar et al., 2010; Röder et al., 2013), coherent motion detection (Bottari et al., 2018; Hadad et al., 2012; Maurer & Lewis, 2017), visual temporal processing (Badde et al., 2020) and visual feature binding (McKyton et al., 2015; Putzar et al., 2007). These visual deficits in congenital cataract-reversal individuals have been attributed to altered neural development due to the absence of early visual experience, as individuals who suffered from developmental cataracts do not typically display a comparable severity of visual impairments (Lewis & Maurer, 2009; Sourav et al., 2020). While the extant literature has reported correlations between structural changes and behavioral outcomes in congenital cataract-reversal individuals (Feng et al., 2021; Guerreiro et al., 2015; Hölig et al., 2023; Pedersini et al., 2023), functional brain imaging (Heitmann et al., 2023; Raczy et al., 2022) and electrophysiological research (Bottari et al., 2016; Ossandón et al., 2023; Pant et al., 2023; Pitchaimuthu et al., 2021) have started to unravel the neural mechanisms which rely on visual experience during early brain development.

Resting-state activity measured via fMRI suggested increased excitation in the visual cortex of congenital cataract-reversal individuals (Raczy et al., 2022): The amplitude of low frequency (<1 Hz) (blood oxygen level-dependent) fluctuations (ALFF) in the visual cortex was increased in congenital cataract-reversal individuals compared to normally sighted controls when they were scanned with their eyes open. Since similar changes were observed in permanently congenitally blind humans, the authors speculated that congenital visual deprivation resulted in an increased E/I ratio of neural circuits due to impaired neural tuning, which was not reinstated after sight restoration (Raczy et al., 2022). Other studies measured resting-state electroencephalogram (EEG) activity and analyzed periodic (alpha oscillations) (Bottari et al., 2016; Ossandón et al., 2023; Pant et al., 2023) as well as aperiodic activity (Ossandón et al., 2023). Both measures pointed towards an higher E/I ratio of visual cortex in congenital cataract-reversal individuals (Ossandón et al., 2023). In recent research, authors have interpreted the slope of the aperiodic component of the EEG power spectral density function as an indirect indication of the relative level of excitation; the flatter the slope, the higher the assumed E/I ratio (R. Gao et al., 2017; Lombardi et al., 2017; McSweeney et al., 2023; Medel et al., 2020; Molina et al., 2020; Muthukumaraswamy & Liley, 2018; Nanda et al., 2023; Schaworonkow & Voytek, 2021). In fact, prospective studies in children have recently reported a flattening of this slope with age, which was interpreted as increasing levels of excitation with age (Favaro et al., 2023; Hill et al., 2022). Ossandón et al. (2023) observed a flatter slope of the aperiodic power spectrum in the high frequency range (20-40 Hz) but a steeper slope of the low frequency range (1-19 Hz) . This pattern was found in both congenital cataract-reversal individuals, as well as in permanently congenitally blind humans. The low frequency range has often been associated with inhibition (Jensen & Mazaheri, 2010; Lozano-Soldevilla, 2018; Lozano-Soldevilla et al., 2014). However, it has remained unclear how to reconcile EEG resting-state findings for lower and higher frequency ranges.

Two studies with permanently congenitally blind humans employed Magnetic Resonance Spectroscopy (MRS) to investigate the concentration of both, the inhibitory neurotransmitter Gamma-Aminobutyric Acid (GABA) and the excitatory neurotransmitters Glutamate/Glutamine (Glx) as proxy measures of visual cortex inhibition and excitation, respectively (Coullon et al., 2015; Weaver et al., 2013). Glutamate/Glutamine concentration was significantly increased in the “visual” cortex of anophthalmic (n = 5) compared to normally sighted individuals, suggesting increased excitability (Coullon et al., 2015). Preliminary evidence in congenitally permanently blind individuals (n = 9) suggested a decreased GABA concentration in the visual cortex compared to normally sighted individuals (Weaver et al., 2013). Thus, these MRS studies corroborated the hypothesis that a lack of visual input at birth enhances relative excitation in visual cortex compared to typical brain development. However, the degree to which neurotransmitter levels recover following sight restoration after a phase of congenital blindness, and how they related to electrophysiological activity, remained unclear.

Here, we filled this gap: we assessed Glutamate/Glutamine (Glx) and Gamma Aminobutyric Acid (GABA+) concentrations using the MEGA-PRESS sequence (Mescher et al., 1998) in individuals whose sight had been restored, on average, after 12 years of congenital blindness. The ratio of Glx/GABA+ concentration was used as a proxy for the ratio of excitatory to inhibitory neurotransmission (Y. Gao et al., 2024; Grent-’t-Jong et al., 2022; Liu et al., 2015; Narayan et al., 2022; Steel et al., 2020; Takei et al., 2016; L. Zhang et al., 2020). Ten congenital cataract-reversal individuals were compared to age-matched, normally sighted controls at rest. In addition to MRS, EEG was recorded to assess and compare aperiodic activity in the same participants. Participants were tested with eyes open and with eyes closed (MRS and EEG), and while viewing visual stimuli (EEG) which changed in luminance (Pant et al., 2023), since both neurotransmitter levels (Kurcyus et al., 2018) and EEG aperiodic activity (Ossandón et al., 2023) systematically varies between these conditions. We predicted an altered visual cortex Glx/GABA+ concentration ratio in the edited MRS signal in congenital cataract-reversal individuals. Since the aperiodic intercept has been linked to broad band neuronal firing (Manning et al., 2009; Musall et al., 2014; Winawer et al., 2013) and based on prior findings suggesting higher excitation in congenital cataract-reversal individuals (Ossandón et al., 2023; Raczy et al., 2022), we predicted a higher intercept as well as an altered slope of the EEG aperiodic component in this group. We further hypothesized that neurotransmitter changes would be concurrent with changes in the slope and intercept of the EEG aperiodic activity in congenital cataract-reversal individuals (Ossandón et al., 2023). Finally, we exploratorily assessed the relationship between the MRS and EEG parameters, as well as their possible link to visual deprivation history and visual acuity in congenital cataract-reversal individuals.

## METHODS

### Participants

We tested two groups of participants. The first group consisted of 10 individuals with a history of dense bilateral congenital cataracts (CC group, 1 female, Mean Age = 25.8 years, Range = 11 – 43.5). Participants in this group were all recruited at the LV Prasad Eye Institute (Hyderabad, India) and the presence of dense bilateral cataracts at birth was confirmed by ophthalmologists and optometrists based on a combination of the following criteria: clinical diagnosis of bilateral congenital cataract, drawing of the pre-surgery cataract, occlusion of the fundus, nystagmus (a typical consequence of congenital visual deprivation), a family history of bilateral congenital cataracts and a visual acuity of fixating and following light (FFL+) or less prior to surgery, barring cases of absorbed lenses. Absorbed lenses occur specifically in individuals with dense congenital cataracts (Ehrlich, 1948), and were diagnosed based on the morphology of the lens, anterior capsule wrinkling, and plaque or thickness of stroma. Prior to cataract surgery, the intactness of the retina is typically checked. Thus, we can exclude major retinal damage as source of group differences.

Duration of deprivation was calculated as the age of the participant when cataract removal surgery was performed on the first eye. Two participants were operated within the first year of life (at 3 months and 9 months of age), all other participants underwent cataract removal surgery after the age of 6 years (Mean Age at Surgery = 11.8 years, SD = 9.7, Range = 0.2 – 31.4). All participants were tested at least 1 year after surgery (Mean Time since Surgery = 14 years, SD = 9.1, Range = 1.8 - 30.9) (Table 1). Visual acuity was significantly below typical vision in this group (Table 1, Supplementary Material S1).

**Table 1:** Clinical and demographic information of the participants with a history of dense bilateral congenital cataracts (CC) as well as demographic information and visual acuity of age-matched normally sighted control participants (SC). NA indicates that patient’s data for the field were not available. FFL: Fixating and Following Light; CF: Counting Fingers; PL: Perceiving Light. Duration of visual deprivation was calculated by subtracting the date of birth from the date of surgery on the first eye (and thus corresponds to the age at surgery). Time since surgery was calculated by subtracting the date of surgery on the first eye from the date of testing. Visual acuity on the date of testing was measured binocularly with the Freiburg Vision Test (FrACT) (Bach 1996). The second group comprised of 10 normally sighted individuals (SC group, 8 males, Mean Age = 26.3 years, Range = 12 – 41.8). Participants across the two groups were age matched (t_(9)_ = -0.12, p = 0.91). Congenital cataract-reversal individuals were clinically screened at the LV Prasad Eye Institute.

|  | GENDER | AGE | VISUAL<br>ACUITY ON<br>DATE<br>TESTED<br>(LOGMAR) | COMORBIDITIES |  |  | VISUAL<br>ACUITY PRE<br>SURGERY |  | DURATION OF<br>VISUAL<br>DEPRIVATION<br>(YEARS) | TIME SINCE<br>SURGERY<br>(YEARS) | FAMILY<br>HISTORY |
| --- | --- | --- | --- | --- | --- | --- | --- | --- | --- | --- | --- |
|  |  |  |  | ABSORBED<br>LENSES | STRABI<br>-SMUS | NYSTAGMUS | OD | OS |  |  |  |
| CC1 | Male | 17.0 | 0.17 | No | Yes | Yes | FFL | FFL + | 0.2 | 16.8 | No |
|  |  |  |  |  |  |  | - |  |  |  |  |
| CC2 | Male | 43.5 | 0.9 | Yes | Yes | Yes | 1.18 | 1 | 20.8 | 22.7 | Yes |
| CC3 | Male | 18.7 | 0.9 | Yes | Yes | Yes | 1.48 | 1.77 | 15.6 | 3.1 | No |
| CC4 | Male | 15.2 | 0.62 | Yes | NA | Yes | CF<br>at<br>1.5<br>m | CF at<br>3 m | 7.0 | 8.2 | No |
| CC5 | Male | 32.4 | 0.88 | No | Yes | Yes | NA | NA | 14.0 | 18.4 | Yes |
| CC6 | Male | 23.9 | 0.78 | No | Yes | Yes | NA | NA | 6.0 | 17.9 | Yes |
| CC7 | Male | 13.1 | 0.54 | No | Yes | Yes | PL+ | PL+ | 0.8 | 12.4 | No |
| CC8 | Male | 18.3 | 0.66 | Yes | No | Yes | 1.2 | 1.3 | 16.4 | 1.9 | Yes |
| CC9 | Male | 36.9 | 1.34 | No | NA | Yes | NA | NA | 6.0 | 30.9 | Yes |
| CC10 | Female | 38.8 | 1.04 | Yes | Yes | Yes | 1.48 | 1.48 | 31.4 | 7.4 | Yes |
| SC1 | Male | 17.7 | 0.2 |  |  |  |  |  |  |  |  |
| SC2 | Male | 41.9 | -0.27 |  |  |  |  |  |  |  |  |
| SC3 | Male | 19.5 | -0.25 |  |  |  |  |  |  |  |  |
| SC4 | Male | 16.0 | -0.11 |  |  |  |  |  |  |  |  |
| SC5 | Male | 33.3 | -0.12 |  |  |  |  |  |  |  |  |
| SC6 | Male | 24.0 | -0.16 |  |  |  |  |  |  |  |  |
| SC7 | Male | 12.2 | -0.25 |  |  |  |  |  |  |  |  |
| SC8 | Female | 25.1 | -0.28 |  |  |  |  |  |  |  |  |
| SC9 | Female | 36.0 | -0.21 |  |  |  |  |  |  |  |  |
| SC10 | Male | 37.3 | -0.22 |  |  |  |  |  |  |  |  |

Both groups did not self-report any neurological or psychiatric conditions, nor any medications. Additionally, all participants were screened for MRI exclusion criteria using a standard questionnaire from the radiology department. One additional individual was tested in each group; they were excluded from data analysis as their data files were corrupted due to inappropriate file transfer from the scanner. All participants (as well as legal guardians for minors) gave written and informed consent. This study was conducted after approval from the Local Ethical Commission of the LV Prasad Eye Institute (Hyderabad, India) as well as of the Faculty of Psychology and Human Movement, University of Hamburg (Germany).

### Data Collection and Analysis

The present study consisted of three data acquisition parts on the same day: (1) Magnetic Resonance Spectroscopy (MRS, 45-60 min.); (2) Electroencephalography (EEG, 20 min. plus time for capping); (3) visual acuity assessment (3-5 min.).

### Magnetic Resonance Spectroscopy

Participants underwent MRI and MRS scanning at LUCID Diagnostics in Hyderabad (India) with a 3T GE SIGNA Pioneer MRI machine employing a 24-channel head coil. An attendant was present in the scanning room for the duration of the scan to ensure that participants were comfortable and followed the instructions.

A T1 weighted whole brain image was collected for each participant (Repetition Time (TR) = 14.97 ms, Echo Time (TE) = 6.74 ms, Matrix size⍰=⍰512⍰×⍰512, In-plane resolution = 0.43⍰×⍰0.43 mm, Slice thickness = 1.6 mm, Axial slices = 188, Interslice interval= -0.8 mm, Inversion time⍰=⍰500 ms, Flip angle⍰=⍰15°). This structural scan enabled registration of every MRS scan to the participants’ anatomical landmarks (Figure 1). For this scan, participants were instructed to stay as still as possible.

**Figure 1:**
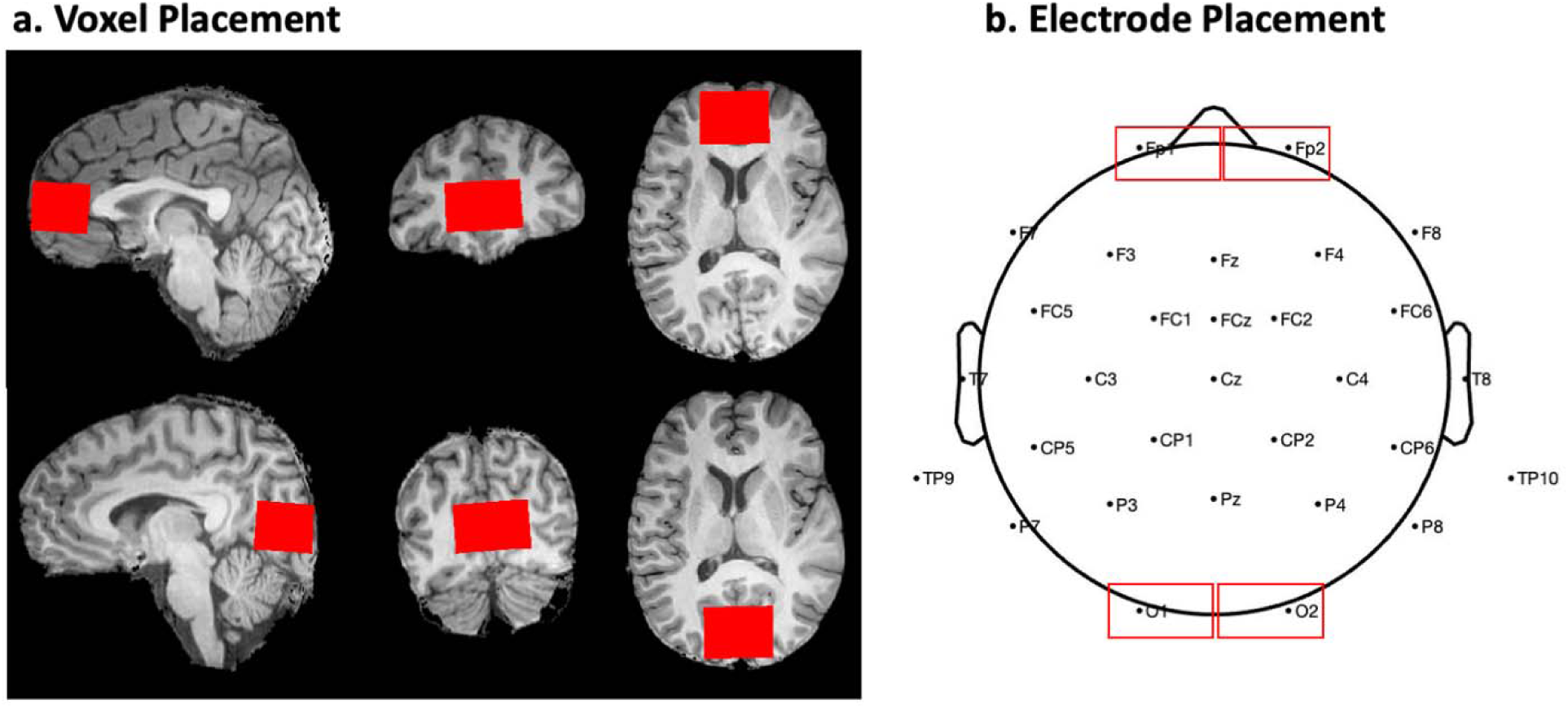
Voxel placement for Magnetic Resonance Spectroscopy and electrode placement for Electroencephalography. a. Position of the frontal cortex (top) and visual cortex (bottom) voxels in a single subject. Skull-stripped figures output from SPM12. b. Electrode montage according to the 10/20 electrode system with marked occipital electrodes preselected for analyses, and frontal electrodes used for control analyses.

The MRS scans consisted of single-voxel spectroscopy data that were collected using the MEGA-PRESS sequence, which allows for in-vivo quantification of the low-concentration metabolites GABA and glutamate+glutamine (Glu+Gln) (Mescher et al., 1998; Mullins et al., 2014). Due to the spectral overlap of GABA (3.0 ppm) and Glu/Gln (3.75 ppm) with the higher concentration peaks of N-Acetyl Aspartate (NAA) and Creatine (Cr), accurate quantification of GABA and Glu/Gln is challenging. MEGA-PRESS uses spectral editing to obtain these measurements. Spectroscopy data consisted of an edit-ON and an edit-OFF spectrum for each voxel, wherein the “ON” and “OFF” refer to whether the frequency of the editing pulse applied is on- or off-resonance with the signal coupled to the GABA complex (applied at approximately 1.9 ppm). Therefore, subtracting repeated acquisitions of the edit-ON and edit-OFF spectra allows for measurement of the magnitude of signals differing in their response to the editing pulse (e.g. GABA), while cancelling out signals that do not (e.g. Cr) (Mescher et al., 1998). Each MEGA-PRESS scan lasted for 8.5 minutes and was acquired with the following specifications: TR = 20001ms, TE = 681ms, Voxel size1=140 mm x 30 mm x 25mm, 256 averages. Additionally, 8 unsuppressed water averages were acquired, allowing all metabolites to be referenced to the tissue water concentration. Concentrations of GABA and Glu/Gln quantified from these acquisitions are respectively referred to as GABA+, due to the presence of macromolecular contaminants in the signal (Mullins et al., 2014), and Glx, due to the combined quantification of the Glu, Gln and Glutathione peaks.

Two MRS scans were collected from the visual cortex, centered on the calcarine sulcus of every participant (Figure 1). A prior study with normally sighted individuals suggested that visual cortex Glx and GABA+ concentrations depend on whether the participants were scanned with eyes open or eyes closed (Kurcyus et al., 2018). Therefore, to ensure any group differences were not potentially driven by differences in eye opening/closure, we tested all participants at rest in two conditions – with eyes open (EO) and eyes closed (EC). Both scans were conducted with regular room illumination, that is, without any explicit visual stimulation.

To ensure that we were identifying neurochemical changes specific to visual regions, we selected the frontal cortex as a control region (Figure 1) and collected two scans (EO and EC) from the frontal cortex. The order of the MRS scans was counterbalanced across individuals for both locations and conditions. Two SC subjects did not complete the frontal cortex scan for the EO condition and were excluded from the statistical comparisons of frontal cortex neurotransmitter concentrations. Voxel placement was optimized to avoid the inclusion of the meninges, ventricles, skull and subcortical structures. For each participant a proper placement was ensured by examining the voxel region across the slices in the acquired T1 volume. Saturation bands to nullify the skull signal were placed at the posterior and anterior edge of the visual cortex and frontal cortex voxel, respectively. Due to the limitations of the clinical scanner settings, rotated and skewed voxels were not possible, and therefore voxels were not always located precisely parallel to the calcarine. As documented in Supplementary Material S2, the visual cortex voxel showed significant (>60%) overlap with the V1-V6 region in every individual participant.

### MRS Data Analysis

All data analyses were performed in MATLAB (R2018b, The MathWorks Inc.). For MRS data analyses we used Gannet 3.0, a MATLAB based toolbox specialized for the quantification of GABA+ and Glx from edited spectrum data (Edden et al., 2014). Following initial data analysis, all datasets were reanalyzed for quantification of NAA, GABA+ and Glx using linear combination modelling with the Osprey toolbox (v. 2.5.0) (Oeltzschner et al., 2020) in MATLAB 2024a (Supplementary Material S3). Osprey had not been released when the study was originally conceptualized. The results did not differ between analysis toolboxes. Here, we present the originally planned analyses with Gannet 3.0.

GABA+ and edited Glx concentration values were obtained and corrected using the *GannetFit, GannetCoRegister,*G*annetSegment* and *GannetQuantify* functions (Edden et al., 2014). Briefly, the reported water-normalized, alpha-corrected concentration values, were corrected for the differences in GABA concentration and relaxation times between different tissue types in the voxel (grey matter, white matter and cerebrospinal fluid) (Harris et al., 2015). Gannet uses SPM12 to determine the proportion of grey matter, white matter and cerebrospinal fluid in each individual participant’s voxel (Penny et al., 2007). Note that the tissue fraction values did not differ between groups or conditions (all p’s > 0.19, see Supplementary Material S4). GABA+, Glx and Glx/GABA+ values were compared across groups as proxy measures of inhibition, excitation and E/I ratio, respectively. The use of Glx/GABA+ as a proxy measure of E/I neurotransmission is supported by a study that observed a regional balance between Glx and GABA+ at 3T (Steel et al., 2020). Further, the Glx/GABA+ ratio has been employed in prior studies of visual (Takei et al., 2016; L. Zhang et al., 2020), cingulate (Bezalel et al., 2019), frontal (Y. Gao et al., 2024; Liu et al., 2015; Narayan et al., 2022) and auditory cortex (Grent-’t-Jong et al., 2022).

To control for potential unspecified visual cortex changes due to eye pathology, as opposed to genuine changes in neurotransmitter ratio, we compared N-Acetyl Aspartate (NAA) concentrations in the visual cortex of CC vs SC individuals. NAA forms one of the most prominent peaks in the MR spectrum (2.0 ppm chemical shift). NAA has been quantified with high reproducibility in the visual cortex (Brooks et al., 1999) and medial-temporal cortex (Träber et al., 2006) of neuro-typical individuals as well as in various pathologies across visual, frontal and temporal cortex (Paslakis et al., 2014), for example, schizophrenia (Mullins et al., 2003). We did not expect to find differences in NAA concentration between CC and SC individuals as it has not been demonstrated to vary in anophthalmia (Coullon et al., 2015) or permanent early blindness (Weaver et al., 2013) in humans. TARQUIN 4.3.11 was employed to analyze the OFF-spectrum data (Wilson et al., 2011) to assess NAA concentration. FID-A toolbox was used to correct the data for phase errors across acquisitions arising from temporal changes in the magnetic field strength or participant motion (Simpson et al., 2017).

The reported values in the results are water-normalized. All data analyses were repeated with Cr-normalized values from Gannet 3.0, and significant results were replicated (Supplementary Material S5).

### MRS Data Quality

The MRS minimum reporting standards form is found in the Supplementary Excel File 1. Mean Signal-to-Noise Ratio values for GABA+ and Glx in all groups and conditions were above 19 in the visual cortex and above 8 in the frontal cortex (Table 2). A recent study has suggested that an SNR value above 3.8 allows for reliable quantification of GABA+ (Zöllner et al., 2021), in conjunction with considering a given study’s sample size (Mikkelsen et al., 2018). Cramer-Rao Lower Bound (CRLB) values, that is, the theoretical lower limit of estimated error, were 30% or lower for NAA quantification in both groups and conditions (Cavassila et al., 2001). Note that CRLB values above 50% are considered unreliable (Wilson et al., 2019). In all quality metrics for Glx, GABA+ and NAA our dataset showed higher quality for the visual cortex voxel than for the frontal cortex voxel, irrespective of group (Main effect of region: all p’s < 0.004, Supplementary Material S6). Such region effects have repeatedly been reported in the MRS literature. They were attributed to magnetic field distortions (Juchem & Graaf, 2017) resulting from the proximity of the frontal cortex voxel to the sinuses. We chose a frontal control voxel rather than a parietal/sensorimotor control voxel (Coullon et al., 2015; Weaver et al., 2013) due to well documented changes in multisensory cortical region as a consequence of congenital blindness (Harrar et al., 2018; Henschke et al., 2017; F. Jiang et al., 2016; Röder et al., 1999; Sabourin et al., 2022; Zatorre et al., 2012) . The fit error for the frontal cortex voxel was below 8.31% for GABA+ and Glx in both groups (Table 2). No absolute cutoffs exist for fit errors. However, Mikkelsen et al. reported a mean GABA+ fit error of 6.24 +/-1.95% from a posterior cingulate cortex voxel across 8 GE scanners using the Gannet pipeline (Mikkelsen et al., 2017). Previous studies in special populations have used frontal cortex data with a fit error of <10% to identify differences between cohorts (Y. Gao et al., 2024; Maier et al., 2022; Pitchaimuthu et al., 2017). Importantly, in the present study, data quality did not significantly differ between groups for GABA+, Glx or NAA (Supplementary Material S6), making it highly unlikely that data quality differences contributed to group differences.

**Table 2:** Quality metrics for Magnetic Resonance Spectroscopy data. Mean quality metrics in each group are reported with the standard deviation in parentheses. The displayed quality metrics for Signal-to-Noise Ratio, Full-Width Half Maxima and Fit Error are those output by Gannet 3.0: Signal-to-noise-ratio (SNR), was calculated in GannetFit.m by estimating the noise in the GABA+/Glx/NAA signal across acquisitions and by dividing the absolute peak height of the GABA+/Glx/NAA signal by the estimated noise; Full-width-half-maxima (FWHM), is defined as the width of the peak in Hertz (Hz); and Fit Error, is defined as the standard deviation of the residual of the GABA+/Glx/NAA peak fit. The Fit Error is expressed as a percentage of the GABA+/Glx peak height. The Cramer-Rao Lower Bound is reported as output by TARQUIN 4.3.11 for the NAA signal (not calculated for GABA+ Glx as these metabolites were quantified using Gannet 3.0). Prior to in-vivo scanning, we confirmed the GABA+ and GABA+/Glx quantification quality with phantom testing (Henry et al., 2011; Jenkins et al., 2019). Imaging sequences were robust in identifying differences of 0.02 mM in GABA concentration; the known vs. measured concentrations of both GABA (r = 0.81, p = 0.004) and GABA/Glx (r = 0.71, p = 0.019) showed significant agreement. This 0.02 mM difference was documented by Weaver et al. (2013) between the occipital cortices of early blind and sighted individuals. The detailed procedure and results are described in Supplementary Material S7. The spectra from all individual subjects are shown in Supplementary Material S8.

| Signal-to-Noise Ratio |  |  |  |
| --- | --- | --- | --- |
|  |  | CC | SC |
| Visual Cortex | NAA | 293.16 (47.50) | 289.01 (50.91) |
|  | GABA+ | 21.53 (3.66) | 19.08 (3.99) |
|  | Glx | 23.75 (3.75) | 22.18 (5.26) |
| Frontal Cortex | NAA | 108.37 (21.84) | 97.20 (28.08) |
|  | GABA+ | 10.311 (2.20) | 8.30 (1.93) |
|  | Glx | 15.82 (4.85) | 13.58 (3.86) |
| Full-Width-Half Maxima |  |  |  |
|  |  | CC | SC |
| Visual Cortex | NAA | 9.04 (0.94) | 8.69 (0.75) |
|  | GABA+ | 19.84 (1.13) | 19.10 (0.71) |
|  | Glx | 16.62 (1.63) | 16.46 (1.63) |
| Frontal Cortex | NAA | 19.26 (2.33) | 21.42 (3.79) |
|  | GABA+ | 21.69 (3.15) | 23.23 (3.41) |
|  | Glx | 27.54 (8.70) | 30.63 (12.64) |
| **Fit Error** |  |  |  |
|  |  | CC | SC |
| Visual Cortex | NAA | 0.81 (0.20) | 0.77 (0.15) |
|  | GABA+ | 3.42 (0.63) | 3.68 (0.63) |
|  | Glx | 3.10 (0.58) | 3.18 (0.47) |
| Frontal Cortex | NAA | 1.33 (0.41) | 1.70 (0.57) |
|  | GABA+ | 6.57 (2.20) | 8.31 (3.65) |
|  | Glx | 4.44 (1.54) | 5.15 (1.90) |
| **Cramer-Rao Lower Bound** |  |  |  |
|  |  | CC | SC |
| Visual Cortex | NAA | 0.13 (0.02) | 0.14 (0.03) |
| Frontal Cortex | NAA | 0.33 (0.22) | 0.26 (0.24) |

#### MRS Statistical Analysis

All statistical analyses were performed using MATLAB R2018b and R v3.6.3.

We compared the visual cortex concentrations of 3 neurochemicals (GABA+ and Glx from the DIFF spectrum, NAA from the edit-OFF spectrum) between the two groups. For each metabolite we submitted the concentration values from the visual cortices of CC and SC individuals to a group (2 Levels: CC, SC)-by-condition (2 Levels: EO, EC) ANOVA model. To compare the Glx/GABA+ ratio between groups, we additionally submitted this ratio value to a group-by-condition ANOVA. Identical analyses were performed for the corresponding frontal cortex neurotransmitter values. Wherever necessary, post-hoc comparisons were performed using t-tests. The data were tested for normality (Shapiro-Wilk) and homogeneity of variance (Levene’s Test) in R v3.6.3 (Supplementary Material S9). In all ANOVA models, the residuals did not significantly differ from normality.

#### Electrophysiological recordings

EEG data analyzed in the present study are a subset of datasets that were included in previous reports (Ossandón et al., 2023; Pant et al., 2023). The EEG datasets were re-analyzed to investigate aperiodic activity in the same participants who took part in the MRS study. MRS and EEG data were acquired on the same day. The EEG was recorded in three conditions: (1) at rest with eyes open (EO, 3 minutes), (2) at rest with eyes closed (EC, 3 minutes) and (3) during visual stimulation with stimuli that changed in luminance (LU) with equal power at all frequencies (0-30 Hz) (Pant et al., 2023). We used the slope of the aperiodic (1/f) component of the EEG spectrum as an estimate of E/I ratio (R. Gao et al., 2017; Medel et al., 2020; Muthukumaraswamy & Liley, 2018) and the intercept as an estimate of broadband neuronal firing activity (Haller et al., 2018; Manning et al., 2009; Miller, 2010).

The EEG was recorded using Ag/AgCl electrodes attached according the 10/20 system(Homan et al., 1987) to an elastic cap (EASYCAP GmbH, Herrsching, Germany) (Figure 1). We acquired 32 channel EEG using the BrainAmp amplifier, with a bandwidth of 0.01–200⍰Hz, sampling rate of 5⍰kHz and a time constant of 0.016 Hz /(10 s) (http://www.brainproducts.com/). All scalp recordings were performed against a left ear lobe reference. Electrode impedance was kept below 10 kOhm in all participants.

Participants were asked to sit as still as possible while the EEG was recorded. First, resting-state EEG data were collected. During the EO condition, participants were asked to look towards a blank screen and to avoid eye movements. During the EC condition, participants were instructed to keep their eyes closed. The order of conditions was randomized across participants.

Subsequently, EEG data were recorded during 100 trials of a target detection task with stimuli that changed in luminance (LU). Stimuli were presented with a Dell laptop, on a Dell 22-inch LCD monitor with a refresh rate of 60⍰Hz. They were created with MATLAB r2018b (The MathWorks, Inc., Natick, MA) and the Psychtoolbox 3 (Brainard, 1997; Kleiner et al., 2007). On each trial, participants observed a circle at the center of a black screen, subtending a visual angle of 17 degrees. The circle appeared for 6.25 s and changed in luminance with equal power at all frequencies (0-30 Hz). At the end of every trial, participants had to indicate whether a target square, subtending a visual angle of 6 degrees, appeared on that trial. The experiment was performed in a darkened room (for further details, see (Pant et al., 2023).

### EEG Data Analysis

Data analysis was performed using the EEGLab toolbox on MATLAB 2018b (Delorme & Makeig, 2004). All EEG datasets were filtered using a Hamming windowed sinc FIR filter, with a high-pass cutoff at 1 Hz and a low-pass cutoff at 45 Hz. A prior version of the analysis was conducted with line noise removal via spectrum interpolation (Ossandón et al., 2023). However, the analyses reported here did not include this step, since we implemented a low-pass cutoff (20 Hz) which falls far below the typical line noise frequency (50 Hz). Eye movement artifacts were detected in the EEG datasets via independent component analysis using the *runica.m* function’s *Infomax* algorithm in EEGLab. Components corresponding to horizontal or vertical eye movements were identified via visual inspection based on criteria discussed in Plöchl et al (Plöchl et al., 2012) and removed.

The two 3 minutes long resting-state recordings (EC, EO) were divided into epochs of 1 s. Epochs with signals exceeding ±120 μV were rejected for all electrodes (see Supplementary Material S10 for percentages by group and condition). We then calculated the power spectral density of the EO and EC resting-state data using the *pwelch* function (sampling rate = 1000 Hz, window length = 1000 samples, overlap = 0).

Datasets collected while participants viewed visual stimuli that changed in luminance (LU) were downsampled to 60⍰Hz (antialiasing filtering performed by EEGLab’s *pop_resample* function) to match the stimulation rate. The datasets were divided into 6.25 s long epochs corresponding to the duration of visual stimulation per trial. Subsequently, baseline removal was conducted by subtracting the mean activity across the length of an epoch (1 s for the EO and EC conditions, 6.25 s for the LU condition) from every data point. After baseline removal, epochs with signals exceeding a threshold of ±120 μV were rejected in order to exclude potential artifacts. Finally, we calculated the power spectral density of the LU data using the *pwelch* function (sampling rate = 60 Hz, window length = 60 samples, overlap = 0).

We derived the aperiodic (1/f) component of the power spectrum for the EO, EC and LU conditions (Donoghue, Haller, et al., 2020; Schaworonkow & Voytek, 2021). First, we fit the 1/f distribution function to the frequency spectrum of each participant, separately for each electrode. The 1/f distribution was fit to the normalized spectrum converted to log-log scale (range = 1-20 Hz) (Donoghue, Dominguez, et al., 2020; Gyurkovics et al., 2021; Schaworonkow & Voytek, 2021). We excluded the alpha range (8-14 Hz) for this fit to avoid biasing the results due to documented differences in alpha activity between CC and SC individuals (Bottari et al., 2016; Ossandón et al., 2023; Pant et al., 2023). This 1/f fit resulted in a value of the aperiodic slope, an aperiodic intercept value corresponding to the broadband power of 1-20 Hz, and a fit error value for the spectrum of every participant, individually for each electrode. Spectra from individual subjects are displayed in Supplementary Material S11. The visual cortex aperiodic slope and intercept values were obtained by averaging across the pre-selected occipital electrodes O1 and O2, resulting in one value of broadband slope and one value of intercept per participant and condition (Figure 1). This procedure yielded average R^2^ values > 0.91 for the aperiodic fit in each group and condition (Supplementary Material S11).

### EEG Statistical Analysis

We compared the average visual cortex aperiodic slope and intercept in separate group (2 Levels: CC, SC) by condition (3 levels: EC, EO, LU) ANOVA models. The data were tested for normality (Shapiro-Wilk) and homogeneity of variance (Levene’s Test) in R v3.6.3) (see Supplementary Material S9); in all ANOVA models, the residuals did not significantly differ from normality.

#### Visual acuity

Visual acuity was measured binocularly for every participant on the date of testing, using the Freiburg Visual Acuity Test (FrACT) (Bach 1996, Bach 2007, https://michaelbach.de/fract/). Visual acuity is reported as the logarithm of the minimum angle of resolution (logMAR, Table 1), wherein higher values indicate worse vision (Elliott, 2016). Analogous to previous studies, we ran a number of exploratory correlation analyses between GABA+, Glx and Glx/GABA+ concentrations and visual acuity at the date of testing, duration of visual deprivation, and time since surgery, respectively, in the CC group (Birch et al., 2009; Guerreiro et al., 2015; Kalia et al., 2014; Rajendran et al., 2020). As expected from normal vision in the SC group, they did not show considerable variance in visual acuity (Table 1); thus, we refrained from calculation correlations between visual acuity and MRS/EEG parameters in the SC group. Based on the literature, we additionally tested the correlation between the neurotransmitter levels and chronological age across the CC and SC groups. All reported correlation coefficients are Pearson correlations, and 95% confidence intervals were calculated for all correlation coefficients.

#### Exploratory correlation analyses between MRS and EEG measures

Exploratory correlation analyses between EEG and MRS measures were run separately for CC and SC individuals. We calculated Pearson correlations between the aperiodic intercept and GABA+, Glx and Glx/GABA+ concentrations. Further, Pearson correlations between the aperiodic slope, and the concentrations of GABA+, Glx and Glx/GABA+ were assessed. MRS measures collected at rest with eyes open (EO) and eyes closed (EC) were correlated with the corresponding resting-state EEG conditions (EO, EC). EEG metrics for the visual stimulation (LU) condition with flickering stimuli were tested for correlation with GABA+, Glx and Glx/GABA+ concentration measured while participants’ eyes were open at rest. We did not have prior hypotheses as to the best of our knowledge no extant literature has tested the correlation between aperiodic EEG activity and MRS measures of GABA+,Glx and Glx/GABA+. Therefore, we corrected for multiple comparisons using the Bonferroni correction (6 comparisons).

## RESULTS

### Transient congenital visual deprivation lowered the Glx/GABA+ concentration in the visual cortex

The Glx/GABA+ concentration ratio was significantly lower in the visual cortex of congenital cataract-reversal (CC) than age-matched, normally sighted control (SC) individuals (main effect of group: F(1,39) = 5.80, p = 0.021, η_p_² = 0.14) (Figure 2). This effect did not vary with eye closure (main effect of condition: F(1,39) = 2.29, p = 0.139, η_p_² = 0.06, group-by-condition interaction: F(1,39) = 1.15, p = 0.290, η_p_² = 0.03). As a control for unspecific effects of surgery or related to visual deprivation on neurochemistry, the frontal cortex Glx/GABA+ concentration was compared between groups. There was no difference between CC and SC individuals in their frontal cortex Glx/GABA+ concentration (main effect of group: F(1,37) = 0.05, p = 0.82, η_p_² < 0.01, main effect of condition: F(1,37) = 2.98, p = 0.093, η_p_² = 0.07, group-by-condition interaction: F(1,37) = 0.09, p = 0.76, η_p_² < 0.01) (Figure 2).

**Figure 2:**
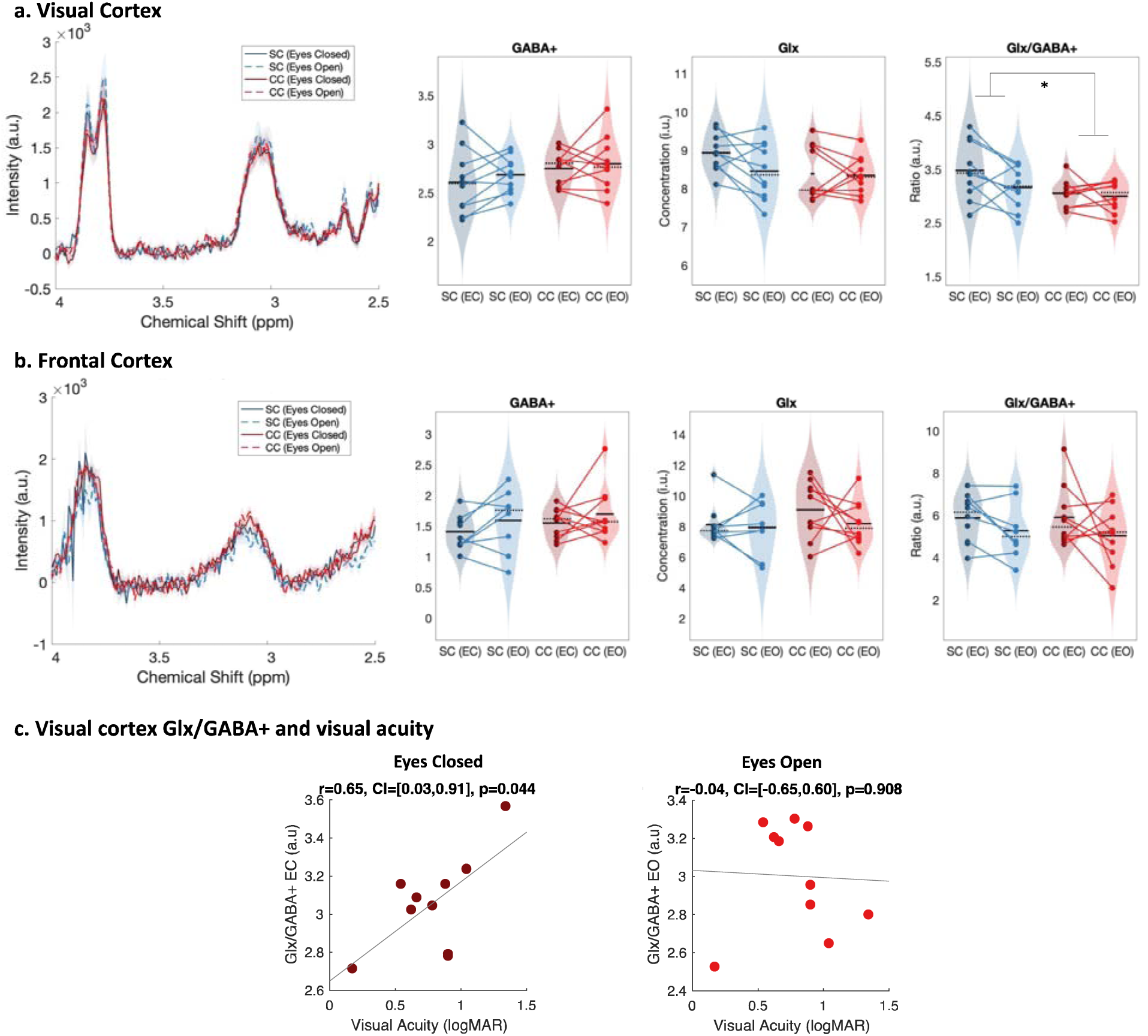
Edited spectra obtained from Magnetic Resonance Spectroscopy (MRS). a. Average edited spectra showing GABA+ and edited Glx peaks in the visual cortices of normally sighted individuals (SC, green) and individuals with reversed congenital cataracts (CC, red) are shown. Edited MRS DIFF spectra are separately displayed for the eyes open (EO), and eyes closed (EC) conditions using dashed and solid lines respectively. The standard error of the mean is shaded. Water-normalized GABA+, water-normalized Glx, and Glx/GABA+ concentration distributions for each group and condition are depicted as violin plots on the right. The solid black lines indicate mean values, and dotted lines indicate median values. The coloured lines connect values of individual participants across conditions. b. Corresponding average edited MRS spectra and water-normalized GABA+, water-normalized Glx and Glx/GABA+ concentration distributions measured from the frontal cortex are displayed. c. Correlations between visual cortex Glx/GABA+ concentrations in the visual cortex of CC individuals and visual acuity in logMAR units are depicted for the eyes closed (EC, left) and eyes open (EO, right) conditions. The 95% confidence intervals (CI) of the correlation coefficients (r) are reported.

When separately comparing CC and SC individuals’ GABA+ and Glx concentrations in the visual cortex, we did not find any significant group difference (GABA+ main effect of group: F(1,39) = 2.5, p = 0.12, η_p_² = 0.06, main effect of condition: F(1,39) = 0.6, p = 0.43, η_p_² = 0.02, group-by-condition interaction: F(1,39) = 0.03, p = 0.86, η_p_² < 0.001; Glx main effect of group: F(1,39) = 2.8, p = 0.103, η_p_² = 0.07, main effect of condition: F(1,39) = 1.8, p = 0.19, η_p_² = 0.05, group-by-condition interaction: F(1,39) = 1.27, p = 0.27, η_p_² -0.03) (Figure 2). In the frontal cortex, GABA+ and Glx concentrations did not vary either with group or condition (all p values > 0.19, all η_p_² < 0.05) (Figure 2). Note that these findings were replicated when Osprey’s quantification method was used: Glx/GABA+ concentration was lower in the visual cortex of CC than SC individuals, while GABA+ and Glx concentration did not significantly differ (Supplementary Material S3). When analyses were repeated with Cr-normalized Glx/GABA+ concentrations, the Glx concentration was found to be significantly lower in CC vs SC individuals’ visual cortices, in addition to the lower Glx/GABA+ concentration ratio (Supplementary Material S5). Since this finding was not replicated with water normalized Glx concentration in Gannet or Osprey, we refrain from interpreting this additional group effect for Glx.

The Glx/GABA+ concentration measured when CC individuals’ eyes were closed correlated positively with visual acuity on the logMAR scale (r = 0.65, p = 0.044), indicating that CC individuals with higher Glx/GABA+ values had worse visual acuity (Figure 2C, Supplementary Material S12). The same correlation was not significant for the eyes opened condition (r = -0.042, p = 0.908) (Figure 2C). Duration of deprivation and time since surgery did not significantly predict Glx/GABA+, GABA+ or Glx concentrations in the CC group (all p values > 0.088, Supplementary Material S12).

No difference in NAA concentration between CC and SC individuals’ visual cortices.

As a control measure to ensure that between-group differences were specific to hypothesized changes in Glx and GABA+ concentrations, we compared the NAA concentration between CC and SC individuals. The NAA concentration did not significantly differ between groups, neither in visual (main effect of group: F(1,39) = 0.03, p = 0.87, η_p_² < 0.001, main effect of condition: F(1,39) = 0.31, p = 0.58, η_p_² <0.01, group-by-condition interaction: F(1,39) = 0.09, p = 0.76, η_p_² <0.01) nor frontal cortex (main effect of group: F(1,37) = 1.1, p = 0.297, η_p_² = 0.02, main effect of condition: F(1,37) = 0.14, p = 0.71, η_p_² = 0.01, group-by-condition interaction: F(1,37) = 0.03, p = 0.86, η_p_² < 0.001) (Supplementary Material S13, Figure S13).

Transient congenital visual deprivation resulted in a steeper aperiodic slope and higher aperiodic intercept at occipital sites.

The aperiodic slope (1-20 Hz), measured via EEG as an electrophysiological estimate of the E/I ratio (R. Gao et al., 2017; Muthukumaraswamy & Liley, 2018), was compared between CC and SC individuals. The aperiodic slope was significantly steeper i.e. more negative, at occipital electrodes in CC than in SC individuals (F(1,59) = 13.1, p < 0.001, η_p_² = 0.19) (Figure 3). Eye closure and visual stimulation did not affect the steepness of the aperiodic slope (F(2,59) = 0.78, p = 0.465, η_p_² = 0.03, group-by-condition interaction: F(2,59) = 0.12, p = 0.885, η_p_² < 0.01).

**Figure 3:**
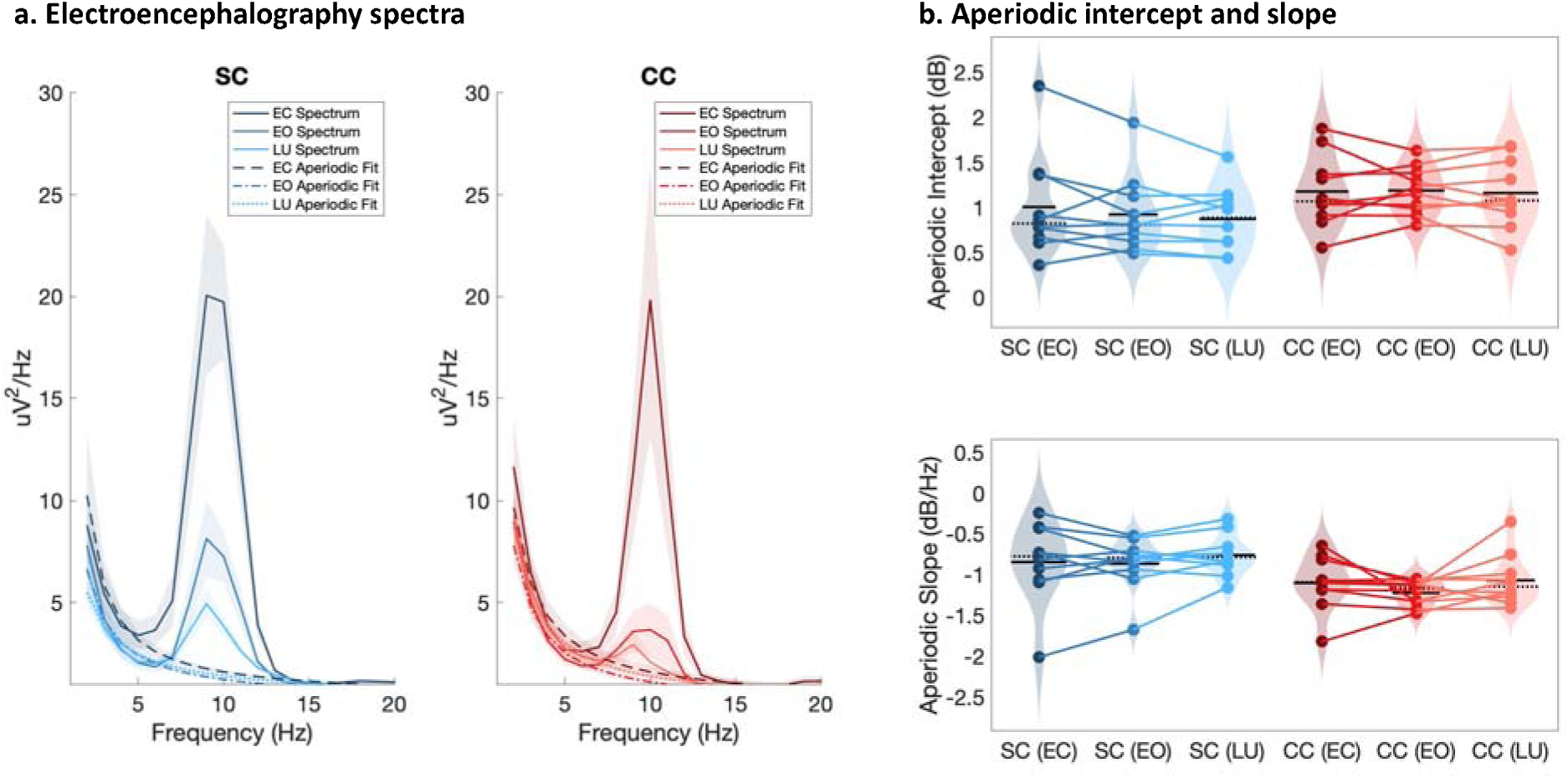
Full spectrum and aperiodic activity of the electroencephalogram (EEG). a. EEG spectra across O1 and O2 with the corresponding aperiodic (1/f) fits for normally sighted individuals (SC, blue, left) and individuals with reversed congenital cataracts (CC, red, right). Spectra of EEG recordings are displayed for the eyes closed (EC) and eyes opened (EO) conditions, as well as while viewing stimuli that changed in luminance (LU). Shaded regions represent the standard error of the mean. b. Aperiodic intercept (top) and slope (bottom) value distributions for each group and condition are displayed as violin plots. Solid black lines indicate mean values, dotted black lines indicate median values. Coloured lines connect values of individual participants across conditions.

The aperiodic intercept (1-20 Hz) was compared between CC and SC individual to estimate group differences in broadband neural activity (Manning et al., 2009; Musall et al., 2014; Winawer et al., 2013) and was found to be significantly larger at occipital electrodes in CC than SC individuals (main effect of group: F(1,59) = 5.2, p = 0.026, η_p_² = 0.09) (Figure 3). Eye closure did not affect the magnitude of the aperiodic intercept in either group (main effect of condition: F(2,59) = 0.16, p = 0.848, η_p_² < 0.01, group-by-condition interaction: F(2,59) = 0.11, p = 0.892, η_p_² < 0.01). No significant group differences in slope and intercept were found for frontal electrodes (Supplementary Material S14).

Within the CC group, visual acuity, time since surgery and duration of blindness did not significantly correlate with the aperiodic slope or the intercept (all p’s > 0.083, Supplementary Material S15). Age negatively correlated with the aperiodic intercept across CC and SC individuals, that is, a reduction of the intercept was observed with age. Similar effects of chronological age have been previously observed (Hill et al., 2022; Voytek et al., 2015) (Supplementary Material S15).

Glx concentration predicted the aperiodic intercept in CC individuals’ visual cortices during ambient and flickering visual stimulation.

We exploratorily tested the relationship between Glx, GABA+ and Glx/GABA+ measured at rest and the EEG aperiodic intercept measured at rest and during flickering visual stimulation, separately for the CC and the SC group. Visual cortex Glx concentration in CC individuals was positively correlated with the aperiodic intercept either when participants had their eyes open during rest (r = 0.91, p = 0.001, Bonferroni corrected) or when they viewed flickering stimuli (r = 0.90, p < 0.001, Bonferroni corrected). Corresponding correlations were not significant for Glx concentrations in the eyes closed condition ( r = 0.341, p > 0.99, Bonferroni corrected). Moreover, in SC individuals no significant correlation was observed between visual cortex Glx concentration and aperiodic intercept in any condition (all p’s > 0.99, Bonferroni corrected) (Figure 4). Given the correlation between the aperiodic intercept and chronological age across groups (Supplementary Material S15), we performed a post-hoc linear regression analysis to model the aperiodic intercept in the CC group with both age and Glx concentration as covariates. Glx concentration, but not age, significantly predicted the aperiodic intercept within the CC group during rest with eyes open and during visual stimulation (Supplementary Material S16).

**Figure 4:**
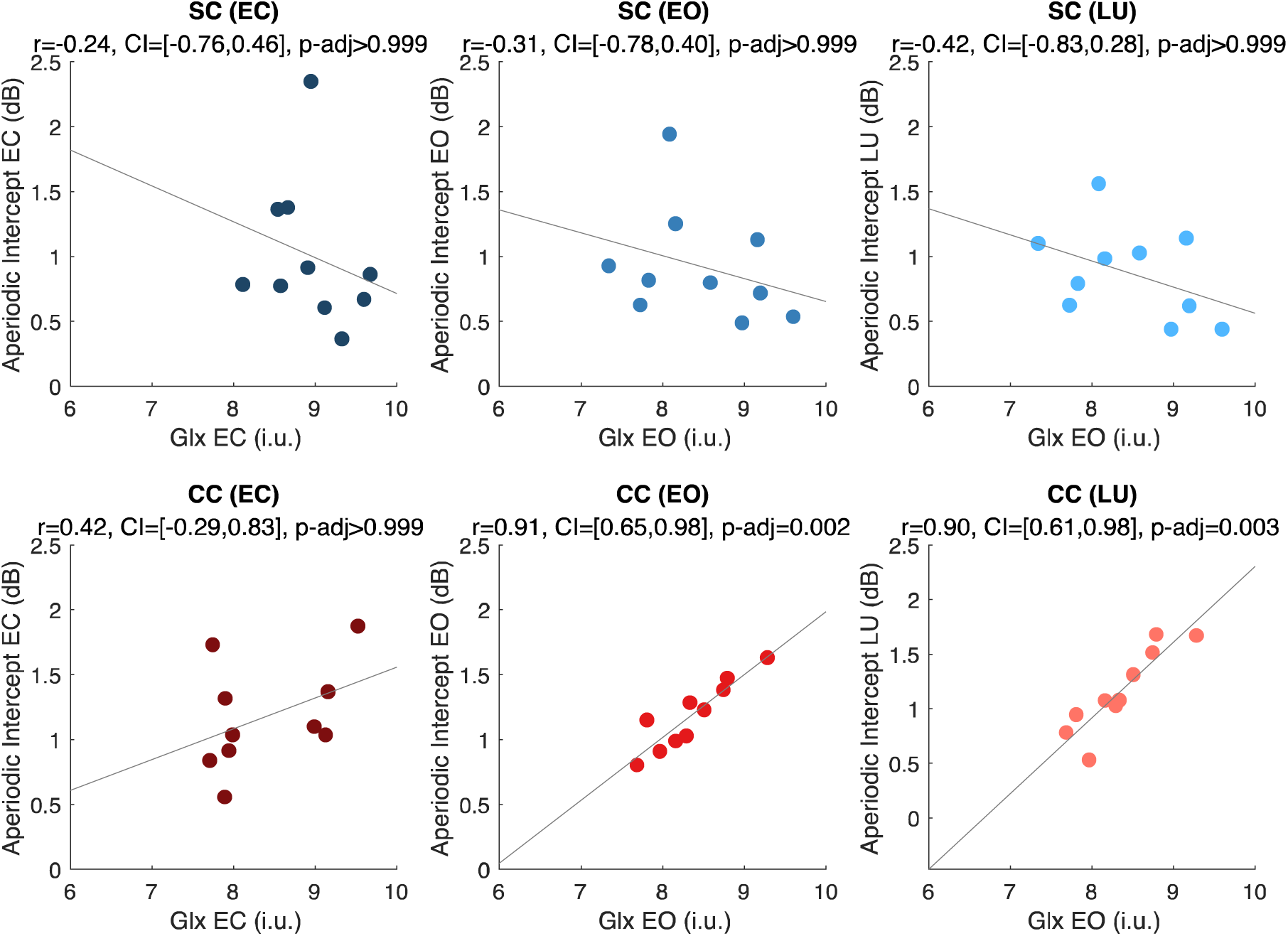
Exploratory correlation analyses between the aperiodic intercept (1-20 Hz) and glutamate/glutamine (Glx) concentration in the visual cortex. Correlations between water-normalized Glx concentration and aperiodic intercept are shown for the eyes closed (EC, left), eyes open (EO, middle) and visual stimulation (LU, right) conditions for sighted controls (SC, green, top) and individuals with reversed congenital cataracts (CC, red, bottom). The reported adjusted p values (p-adj) are Bonferroni corrected for multiple comparisons. The 95% confidence intervals (CI) of the correlation coefficients (r) are reported.

A negative correlation between the aperiodic slope and Glx concentration in CC individuals (i.e., steeper slopes with increasing Glx concentration) was observed during visual stimulation, but did not survive correction for multiple comparisons (Supplementary Material S17). No such correlation was observed between Glx concentration and aperiodic slope in the eyes open or closed conditions. Visual cortex GABA+ concentration and Glx/GABA+ concentration ratios did not significantly correlate with the aperiodic intercept or slope in either CC or SC individuals, during any experimental condition (Supplementary Material S17).

## DISCUSSION

Research in non-human animals has provided convincing evidence that the ratio of excitation to inhibition (E/I) in the visual cortex is reliant on early visual experience (Froemke, 2015; Haider et al., 2006; Hensch et al., 1998; Takesian & Hensch, 2013; Wu et al., 2022). Studies in humans who were born blind due to dense bilateral cataracts, and who had received sight restoration surgery in childhood or as adults, have found limited recovery of both basic visual and higher order visual functions (Birch et al., 2009; Röder & Kekunnaya, 2021). Here, we tested whether neurotransmitter concentrations and electrophysiological markers of cortical E/I ratio depend on early visual experience in humans, and how possible changes in visual cortex E/I ratio relate to sight recovery. First, we employed Magnetic Resonance Spectroscopy (MRS) and assessed Glutamate/Glutamine (Glx) and Gamma-Aminobutyric Acid (GABA+) concentrations, as well as their ratio, in the visual cortex (Shibata et al., 2017; Steel et al., 2020; Takei et al., 2016). Second, the slope and intercept of the aperiodic resting-state EEG activity with eyes open and eyes closed (R. Gao et al., 2017; Muthukumaraswamy & Liley, 2018; Ossandón et al., 2023), as well as during flickering visual stimulation (Pant et al., 2023), were measured over the occipital cortex in the same individuals. The EEG measures allowed us to exploratorily relate neurotransmitter changes to neural activity changes in congenital cataract-reversal individuals.

We found a lower Glx/GABA+ concentration ratio in the visual cortex of congenital cataract-reversal (CC) individuals as compared to normally sighted controls (SC). Additionally, the slope of the aperiodic EEG power spectrum was steeper for the low frequency range (1-20 Hz) and its intercept was higher in CC than SC individuals. In the CC group, Glx concentration correlated with the intercept of the aperiodic component during flickering visual stimulation. The Glx/GABA+ concentration ratio during the eyes closed condition predicted visual acuity of CC individuals. Together, the present results provide initial evidence for experience-dependent development of the E/I ratio in the human visual cortex, with consequences for behavior.

Previous MRS studies in the visual cortex of permanently congenitally blind humans reported higher Glx concentrations (Coullon et al., 2015) in five anophthalmic humans, and numerically lower GABA concentrations in congenitally blind humans (Weaver et al., 2013) (n = 9), as compared to normally sighted individuals. These results were interpreted as suggesting a higher E/I ratio in the visual cortex of permanently congenitally blind humans, which would be consistent with the extant literature on higher BOLD activity in the visual cortices of the same population (Bedny, 2017; Röder & Kekunnaya, 2022). We observed a lower Glx/GABA+ ratio and a steeper slope of the aperiodic EEG activity (1-20 Hz) at occipital electrodes, both of which suggest a lower rather than higher E/I ratio in the visual cortex of CC individuals. Here, we speculate that our results imply a change in neurotransmitter concentrations as a consequence of *restoring* vision following congenital blindness. Further, we hypothesize that due to limited structural plasticity after a phase of congenital blindness, the neural circuits of CC individuals, which had adapted to blindness after birth, likely employ physiological plasticity mechanisms (Knudsen, 1998; Mower et al., 1985; Röder et al., 2021), in order to re-adapt to the newly available visual excitation following sight restoration later in life.

Structural remodeling (Bourgeois, 1996) for typical E/I balance requires visual experience following birth (Hensch & Fagiolini, 2005; Takesian & Hensch, 2013; H. Zhang et al., 2018) and is linked to a sensitive period (Desai et al., 2002; Hensch & Fagiolini, 2005). A repeatedly documented finding in permanently congenitally blind humans is the increased thickness of visual cortex (Anurova et al., 2015; Hölig et al., 2023; J. Jiang et al., 2009). These structural changes in permanently congenitally blind individuals were interpreted as a lack of experience-dependent pruning of exuberant synapses and/or reduced myelination, the latter typically leading to a shift of the grey-white matter boundary (Natu et al., 2019). In parallel, it was observed in non-human primates that the overproduction of synapses during the initial phase of brain development was independent of experience, but that synaptic pruning, predominantly of excitatory synapses, depended on visual experience (Bourgeois, 1996; Bourgeois et al., 1989). The lack of excitatory synapse pruning was thought to underlie the observed higher excitability of visual cortex due to congenital visual deprivation (Benevento et al., 1992; Huang et al., 2015; Morales et al., 2002). Crucially, increased visual cortex thickness (Feng et al., 2021; Guerreiro et al., 2015; Hölig et al., 2023) and higher BOLD activity during rest with the eyes open (Raczy et al., 2022) have been observed for CC individuals following congenital blindness, suggesting incomplete recovery of cortical structure and function *after* sight restoration in humans. Thus, the restored feedforward drive to visual cortex after cataract removal surgery might reach a visual cortex with a lower threshold for excitation.

As to the best of our knowledge, the present study was the first assessing MRS markers of cortical excitation in humans following sight restoration after congenital blindness, we allude to non-human animal work for the interpretation of our findings. Such studies have often demonstrated that excitation and inhibition go hand-in-hand (Froemke, 2015; Haider et al., 2006; Isaacson & Scanziani, 2011; Tao & Poo, 2005). Analogously, we speculate that an overall reduction in Glx/GABA ratio might be effective in counteracting the aforementioned adaptations to congenital blindness, i.e. a lower threshold for excitation. Higher overall visual cortex excitation as a consequence of congenital blindness (Benevento et al., 1992; Morales et al., 2002) might come with the risk of runaway excitation in the presence of restored visually-elicited excitation. Phrased differently, we postulate that significantly lowering overall excitation of an originally hyper-excited visual cortex (during the phase of blindness) after sight restoration might be instrumental for maintaining neural circuit stability. Evidence for such homeostatic adjustments comes from studies with normally sighted humans that observed a reduction of GABA concentrations in visual cortex (Lunghi et al., 2015) and an increase in the BOLD response (Binda et al., 2018) following monocular blindfolding. Further, studies in adult mice have provided support for a homeostatic adjustment of the E/I ratio following prolonged changes in neural activity (Chen et al., 2022; Goel & Lee, 2007; Keck et al., 2017; Whitt et al., 2013). For example, a long period of decreased activity following enucleation in adult mice commensurately decreased inhibitory drive (Keck et al., 2011), primarily onto excitatory neurons (Barnes et al., 2015). In line with the lowered Glx/GABA+ ratio being a compensatory measure to prevent runaway excitation during visual stimulation, the link between the Glx/GABA+ ratio during eye closure and visual acuity in an exploratory correlation analysis suggests that the more successful such assumed downregulation of the E/I ratio in visual cortex, the better the visual recovery. In fact, this correlation with visual acuity recovery is reminiscent of a previously reported correlation in a larger group of CC individuals, between decreased visual cortex thickness and better visual acuity (Hölig et al., 2023). Hence, CC individuals with more advanced structural normalization appear to have a better starting point for functional recovery, the latter possibly mediated by physiological plasticity. Yet, future work has to explicitly test these hypotheses.

An increased intercept of the aperiodic component of occipital EEG activity was observed in the same CC individuals who underwent MRS assessment, irrespective of condition, that is, during rest with eyes open and eyes closed, as well as during flickering stimulation. The intercept of the aperiodic component has been linked to overall neuronal spiking activity (Manning et al., 2009; Musall et al., 2014) and fMRI BOLD activity (Winawer et al., 2013). The higher aperiodic intercept may therefore signal increased spontaneous spiking activity in the visual cortex of CC individuals. This interpretation would be consistent with the previously observed increase in visual cortex BOLD activity of CC compared to SC individuals (Raczy et al., 2022).

In CC individuals, the intercept of the aperiodic activity was highly correlated with the Glx concentration during rest with eyes open and during flickering stimulation. This exploratory finding needs replication in a larger sample. If reliable, the correlation between the EEG aperiodic intercept and Glx concentration in CC individuals might indicate more broadband firing (Manning et al., 2009; Winawer et al., 2013) in CC than SC individuals during active and passive visual stimulation.

### Limitations

The sample size of the present study was rather high for rare population of carefully diagnosed CC individuals, but undoubtedly overall small. Access to CC individuals was limited by the constraints of the COVID-19 pandemic. Hence, all the group differences, the exploratory correlations with visual history metrics, and between MRS-EEG parameters, are reported for further investigation in a larger sample. Moreover, our speculative accounts for the present findings need to be validated with pre- and post-surgery assessments. Finally, a comparison of CC individuals with a control group of developmental cataract-reversal individuals would be instrumental to test the hypothesis that the observed group differences are specific to early brain development.

We are aware that MRS and EEG has a low spatial specificity. Moreover, MRS measures do not allow us to distinguish between presynaptic, postsynaptic and vesicular neurotransmitter concentrations. However, all reported group differences in MRS and EEG parameters were specific to visual cortex and were not found for the frontal control voxel or at frontal electrodes, respectively. While data quality was lower for the frontal compared to the visual cortex voxels, as has been observed previously (Juchem & Graaf, 2017; Rideaux et al., 2022), this was not an issue for the EEG recordings. Thus, lower sensitivity of frontal measures cannot easily explain the lack of group differences for frontal measures. Crucially, data quality did not differ between groups.

While interpretations of new data in the absence of similar data sets are necessarily speculative, the validity of the neurochemical findings was supported by quality assessments; phantom testing showed high correlations between the experimentally varied metabolite concentrations and the extracted GABA+ and Glx concentrations (Supplementary Figure S3). The neurochemical results were robust to analysis pipelines (Supplementary Material S3) as well as normalization method (Supplementary Material S5). The EEG results from the present group of CC individuals replicated effects observed in a larger sample of 28 additional CC individuals (Supplementary Material S18) (Ossandón et al., 2023), as well as prior findings from another sample reporting lower alpha power (Supplementary Material S19) (Bottari et al., 2016; Pant et al., 2023). Further, the aperiodic intercept of EEG activity decreased with chronological age irrespective of group or condition, replicating earlier reports (Hill et al., 2022; Voytek et al., 2015) (Supplementary Material S15). Finally, group differences were observed despite the considerable variance of blindness duration and time since surgery, demonstrating the crucial role of early visual experience.

### Conclusion

The present study in sight recovery individuals with a history of congenital blindness indicates that E/I balance is a result of early experience and crucial for human behavior. We provide initial evidence that the E/I ratio in congenital cataract-reversal individuals is altered even years after surgery, which may be due to previous adaptation to congenital blindness.

## AUTHOR CONTRIBUTIONS

RP, KP, JO, JF and BR conceptualized the study. RP, KP and IS collected the data. IS and RK diagnosed, recruited and provided clinical assessments of participants. RP, JO and KP analyzed the data. BR and JF supervised data analysis and methodological decisions. BR, RK and SL provided infrastructure, resources and funding. RP and BR wrote the original draft of the manuscript, and all authors provided edits and reviews on the final draft of the manuscript.

## ACKNOWLEDGEMENTS

We thank the technical staff of the Lucid Medical Diagnostics Center, Banjara Hills, Hyderabad, India, in particular Mr. Balakrishna Vaddepally, for technical assistance during collection of MRS/MRI data. We would like to acknowledge Dr. Suddha Sourav for technical support, and Ms. Prativa Regmi for assistance with phantom testing and data collection. We are grateful to D. Balasubramanian of the L.V. Prasad Eye Institute for initiating and supporting our research. The study was funded by the German Research Foundation (DFG Ro 2625/10-1 and SFB 936-178316478-B11) and Landesforschungsförderung (LFF-FV 6) of the Free and Hanseatic City of Hamburg to Brigitte Röder. Rashi Pant was supported by a PhD student fellowship from the Hector Fellow Academy GmbH.

