## Supplementary Material for "Altered visual cortex excitatory/inhibitory ratio following transient congenital visual deprivation in humans"

**S1. Visual acuity**

The lower visual acuity (evidenced by higher logMAR values) in congenital cataract-reversal individuals was expected from a large number of previous reports (Khanna et al., 2013). Binocular visual acuity values measured on the day of MRS/EEG testing with the Frieburg Visual Acuity test (Bach, 2007) are seen in Figure S1 and reported in Table 1 of the Methods.

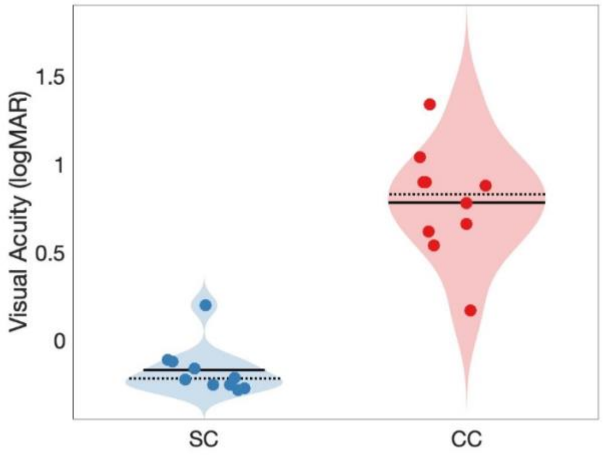


***Figure S1: Visual acuity in normally sighted individuals (SC) and congenital cataract-reversal (CC individuals).*** *Binocularly measured visual acuity distributions in logarithmic of minimum angle of resolution (logMAR) are displayed as violin plots. Solid black lines indicate mean values, dotted black lines indicate median values.*

**S2. Percentage overlap of visual cortex MRS voxel with anatomically defined visual cortex region**

Percentage of overlap between the visual cortex MRS voxel and an anatomically defined visual cortex region of interest (ROI) was calculated for every subject. First, the visual ROI mask was obtained using the Anatomy toolbox in SPM 12 (Eickhoff et al., 2005), including all areas V1-V6 of the occipital lobe. This mask was aligned, co-registered and resliced to every subject's T1 scan. Subsequently, the proportion of vertices in the MRS visual cortex voxel mask (as generated by Gannet 3.0 using SPM 12) overlapping with the T1-aligned occipital lobe mask of each participant was calculated as a percentage of the total number of vertices in the MRS visual cortex voxel. The percentage overlap in both groups did not significantly differ (Mean CC = 67.1%, Mean SC = 70%, $t(18) = -1.14$, $p = 0.269$).

**S3. Magnetic Resonance Spectroscopy data analysis using linear combination modeling (Osprey)**

In the context of MEGA-PRESS data analysis, a recent study suggested that linear combination modeling offers superior reproducibility compared to peak fitting methods, such as using three gaussian peaks (as implemented in Gannet) (Hupfeld et al., 2024). Osprey is an open-source toolbox which uses linear combination modelling for analysis of MEGA-PRESS as well as PRESS datasets (Oeltzschner et al., 2020). As our experiment was conceptualized prior to the release of this toolbox, we originally analyzed our data using Gannet 3.0 by the same authors, and TARQUIN for OFF-spectrum analysis. Subsequently, we re-analyzed our data using Osprey v2.6.0 and found that the main findings of the analysis corresponded to those obtained with Gannet and TARQUIN (Figure S3.1, Figure S3.2). While visual cortex GABA+ (Main effect of group $F(1,39) = 2.48$, $p = 0.124$, $\eta_p^2 = 0.06$, Main effect of condition $F(1,39) = 0.92$, $p = 0.345$, $\eta_p^2 = 0.02$, Group-by-condition interaction $F(1,39) = 0.35$, $p = 0.555$, $\eta_p^2 < 0.01$) and Glx concentration (Main effect of group $F(1,39) = 2.75$, $p = 0.106$, $\eta_p^2 = 0.07$, Main effect of condition $F(1,39) = 0.44$, $p = 0.512$, $\eta_p^2 = 0.01$, Group-by-condition interaction $F(1,39) = 1.46$, $p = 0.234$, $\eta_p^2 = 0.04$) did not significantly differ between CC and SC individuals, the Glx/GABA+ concentration ratio was lower in the visual cortex of CC than SC individuals across conditions (Main effect of group $F(1,39) = 7.67$, $p = 0.009$, $\eta_p^2 = 0.17$, Main effect of condition $F(1,39) = 1.6$, $p = 0.214$, $\eta_p^2 = 0.04$, Group-by-condition interaction $F(1,39) = 0.11$, $p = 0.743$, $\eta_p^2 < 0.01$) (Figure S3.1). NAA concentration did not differ between the visual cortices of CC vs SC individuals, regardless of condition (Main effect of group $F(1,39) = 0.93$, $p = 0.342$, $\eta_p^2 = 0.02$, Main effect of condition $F(1,39) = 0.53$, $p = 0.471$, $\eta_p^2 = 0.01$, Group-by-condition interaction $F(1,39) = 0.14$, $p = 0.714$, $\eta_p^2 < 0.01$) (Figure S3.2).

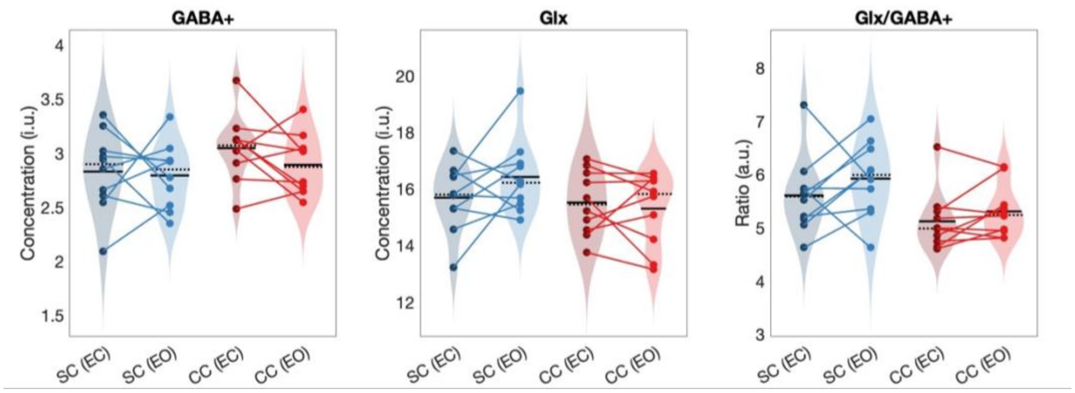


***Figure S3.1: Edited (DIFF) spectrum metabolites quantified via Osprey.*** *Water-normalized and tissue corrected GABA+, water-normalized and tissue-corrected Glx, and Glx/GABA+ concentration distributions from the visual cortex are depicted as violin plots for each group and condition (left to right). The solid black lines indicate mean values, and dotted lines indicate median values. The coloured lines connect*

*values of individual participants across conditions. Results for congenitally cataract-reversal individuals (CC) and for normally sighted controls (SC) are shown in blue and red, respectively. EC=Eyes closed, EO=Eyes open. ,*

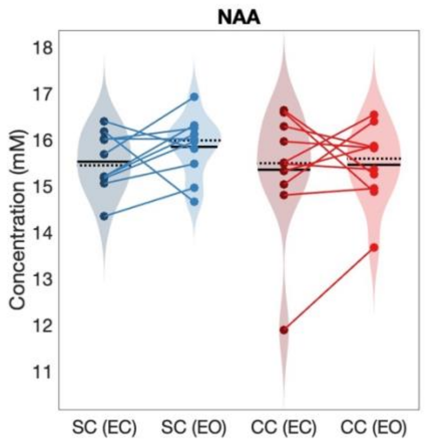


***Figure S3.2: OFF spectrum metabolites quantified via Osprey.*** *Water-normalized NAA concentration distributions from the visual cortex are depicted as violin plots for each group and condition (left to right). The solid black lines indicate mean values, and dotted lines indicate median values. The coloured lines connect values of individual participants across conditions. For abbreviations see Figure S3.1.*

**S4. Tissue Fractions**

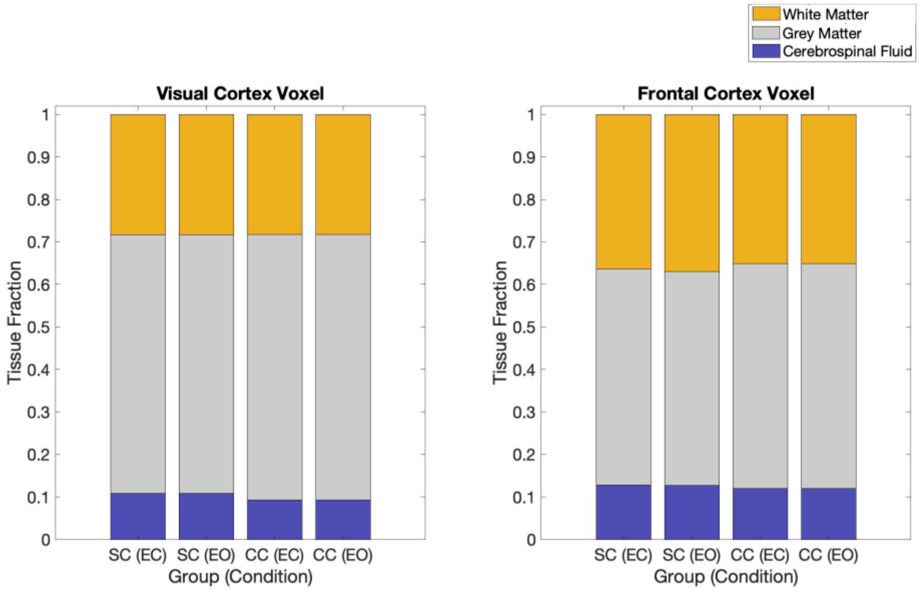


***Figure S2: Tissue fractions for Magnetic Resonance Spectroscopy (MRS) voxels.*** *The fractions of white matter (yellow), grey matter (grey) and cerebrospinal fluid (blue) are displayed for the eyes open (EO), and eyes closed (EC) conditions in the congenital cataract-reversal group (CC) and the normally sighted control group (SC). Tissue fractions were separately calculated for the visual (left) and frontal (right) cortex voxels.*

**S5. Cr-Normalized analysis of Magnetic Resonance Spectroscopy data**

To ensure that our results were not specific to water-normalized quantification of Glx/GABA+, we reran all analyses with the same pipeline specified in the methods section using Creatine (Cr) normalized GABA+ and Glx quantities across the visual cortex of congenital cataract-reversal (CC) and normally sighted control (SC) individuals. Cr is often used as an internal reference as its concentration is relatively stable in most brain regions. Similar to the water-normalized values, a lower Glx/GABA+ concentration ratio was observed in the visual cortex of CC than SC individuals with Cr-normalization (Main effect of group: $F(1,39) = 5.80$, $p = 0.021$, $\eta_p^2 = 0.14$), regardless of eye opening or eye closure (Main effect of condition: $F(1,39) = 2.29$, $p = 0.138$, $\eta_p^2 = 0.06$, Group-by-condition interaction: $F(1,39) = 1.15$, $p = 0.290$, $\eta_p^2 = 0.03$) (Figure S22). Further, Cr-normalized GABA+ concentration did not differ between groups or conditions (Main effect of group: $F(1,39) = 0.82$, $p = 0.369$, $\eta_p^2 = 0.02$, Main effect of condition: $F(1,39) = 0.94$, $p = 0.339$, $\eta_p^2 = 0.02$, Group-by-condition interaction: $F(1,39) = 0.09$, $p = 0.762$, $\eta_p^2 < 0.01$). Notably, unlike water-normalized Glx values (Results, Figure 2), Cr-normalized Glx concentration was lower in the visual cortex of CC than SC individuals (Main effect of group: $F(1,39) = 4.73$, $p = 0.036$, $\eta_p^2 = 0.12$), regardless of condition (Main effect of condition: $F(1,39) = 0.91$, $p = 0.346$, $\eta_p^2 = 0.02$, Group-by-condition interaction: $F(1,39) = 0.94$, $p = 0.339$, $\eta_p^2 = 0.02$) (Figure S5). None of the Cr-normalized values differed by group or condition, nor where there any significant group-by-condition interactions in the corresponding frontal cortex comparison (all $F(1,39) < 2.54$, p's > 0.119, all $\eta_p^2 < 0.07$).

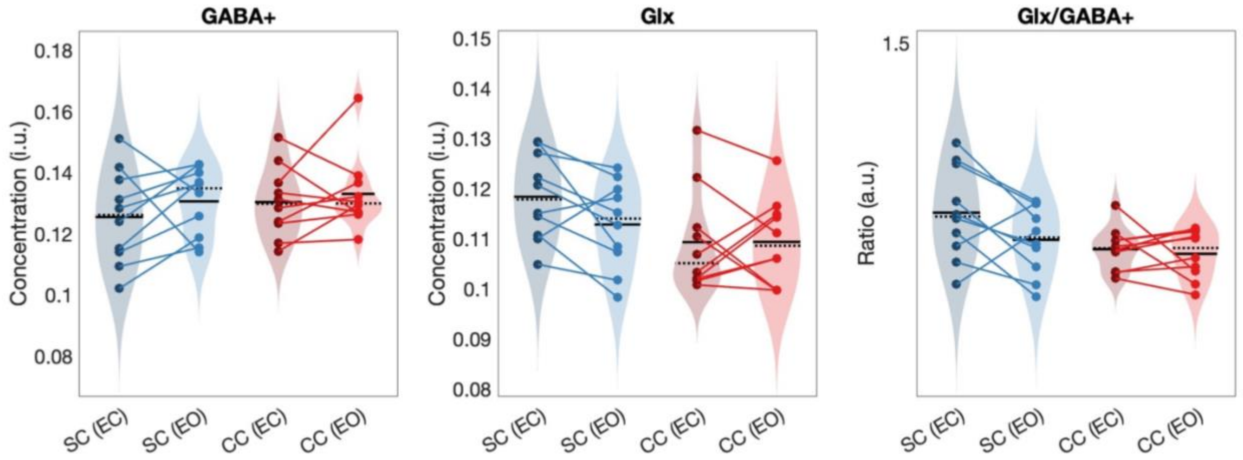


***Figure S5: Cr-normalized edited (DIFF) spectrum metabolites.*** *Creatine (Cr)-normalized GABA+, Cr-normalized Glx, and Glx/GABA+ concentration distributions from the visual cortex are depicted as violin plots for each group and condition (left to right). The solid black lines indicate mean values, and dotted lines indicate median values. The coloured lines connect values of individual participants across conditions.*

**S6. Magnetic Resonance Spectroscopy Quality Metrics analysis**

| | | Main Effect of Group | | | Main Effect of Region | | | Group-by-Region Interaction | | |
|---|---|---|---|---|---|---|---|---|---|---|
| | | F(1,39) | $\eta_p^2$ | p | F(1,39) | $\eta_p^2$ | p | F(1,39) | $\eta_p^2$ | p |
| Signal-to-Noise Ratio | NAA | 0.38 | 0.011 | 0.539 | 232.00 | 0.865 | <0.001 | 0.08 | 0.002 | 0.778 |
| | GABA+ | 3.37 | 0.084 | 0.080 | 127.12 | 0.779 | <0.001 | 0.01 | <0.001 | 0.936 |
| | Glx | 0.39 | 0.011 | 0.534 | 26.75 | 0.426 | <0.001 | <0.001 | <0.001 | 0.989 |
| Full-Width Half Maxima | NAA | 1.53 | 0.041 | 0.224 | 247.71 | 0.873 | <0.001 | 2.94 | 0.076 | 0.095 |
| | GABA+ | 0.09 | 0.002 | 0.765 | 21.71 | 0.376 | <0.001 | 0.56 | 0.015 | 0.457 |
| | Glx | 0.20 | 0.005 | 0.660 | 31.56 | 0.467 | <0.001 | 0.11 | 0.003 | 0.743 |
| Fit Error | NAA | 1.97 | 0.052 | 0.168 | 38.36 | 0.515 | <0.001 | 3.00 | 0.077 | 0.092 |
| | GABA+ | 2.78 | 0.070 | 0.104 | 69.14 | 0.657 | <0.001 | 1.65 | 0.043 | 0.206 |
| | Glx | 0.26 | 0.007 | 0.610 | 12.91 | 0.264 | <0.001 | 0.22 | 0.006 | 0.643 |
| Cramer-Rao Lower Bound | NAA | 0.05 | 0.001 | 0.821 | 9.34 | 0.206 | 0.004 | 0.09 | 0.002 | 0.760 |

***Table S6: ANOVA results for quality metrics on Magnetic Resonance Spectroscopy data.*** *Quality metrics were compared for each signal (GABA+, Glx and NAA) in a group (congenital cataract-reversal, normally sighted control)-by-region (Visual Cortex, Frontal Cortex) ANOVA.*

**S7. Phantom Testing**

Phantom testing was conducted to confirm the quality of the MRS data and analysis pipeline. GABA concentration was varied from 0 to 2 mM in a 1 liter, 7.2 pH phantom with fixed metabolite concentrations of Cr (8 mM), NAA (15 mM), Glutamate (12 mM) and Glutamine (3 mM) (Jenkins et al., 2019). Phosphate Buffer Saline (PBS) was used to maintain the pH at room temperature. The base solution of several metabolites was included to gauge the quality of the overall signal as well as ensuring the similarity of the phantom to metabolites present in-vivo (Jenkins et al., 2019). The range of used GABA concentrations included the previously reported GABA concentration in the visual cortex of congenitally blind individuals (Weaver et al., 2013).

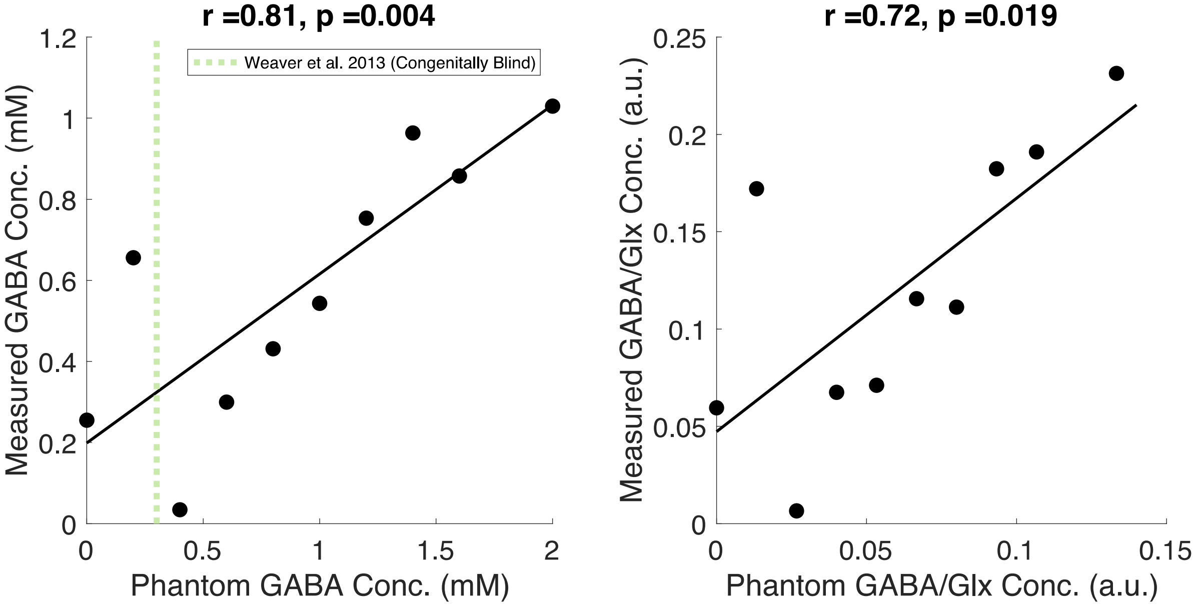


***Figure S7: Phantom testing of GABA concentrations**. Plots depicting the correlation between known and measured concentrations from phantom scans of Gamma-Aminobutyric Acid (GABA; left) and the ratio of GABA to Glutamate/Glutamine (GABA/Glx, right) concentration. In the left panel, previously reported GABA concentration from the visual cortex of congenitally blind individuals* (Weaver et al., 2013) *is marked with a vertical dotted line.*

Eleven phantom scans were obtained varying the known concentration of GABA in steps of 0.2 mM (corresponding to 0.0206 g) (Figure S7), the reported difference in visual cortex GABA concentration between early blind (mean = 0.3 mM) and normally sighted (mean = 0.5 mM) individuals' visual cortex (Weaver et al., 2013). Note that Weaver et al. reported that this group difference did not survive the Bonferroni-Holm correction. Nevertheless, to the best of our knowledge, no other study has reported significant GABA concentration differences between permanently congenitally blind humans and sighted controls based on MRS assessments in humans.

For both GABA and the concentration ratio of GABA/Glx (calculated instead of Glx/GABA due to the 0-GABA concentration solution), our measured values showed significant agreement with the known phantom concentration values (Figure S7). These results demonstrate that our data acquisition and analysis pipeline were adequate to identify differences between CC and SC individuals' visual cortices within previously reported concentration ranges.

### S8. Individual subjects' MRS edited spectra (Visual Cortex)

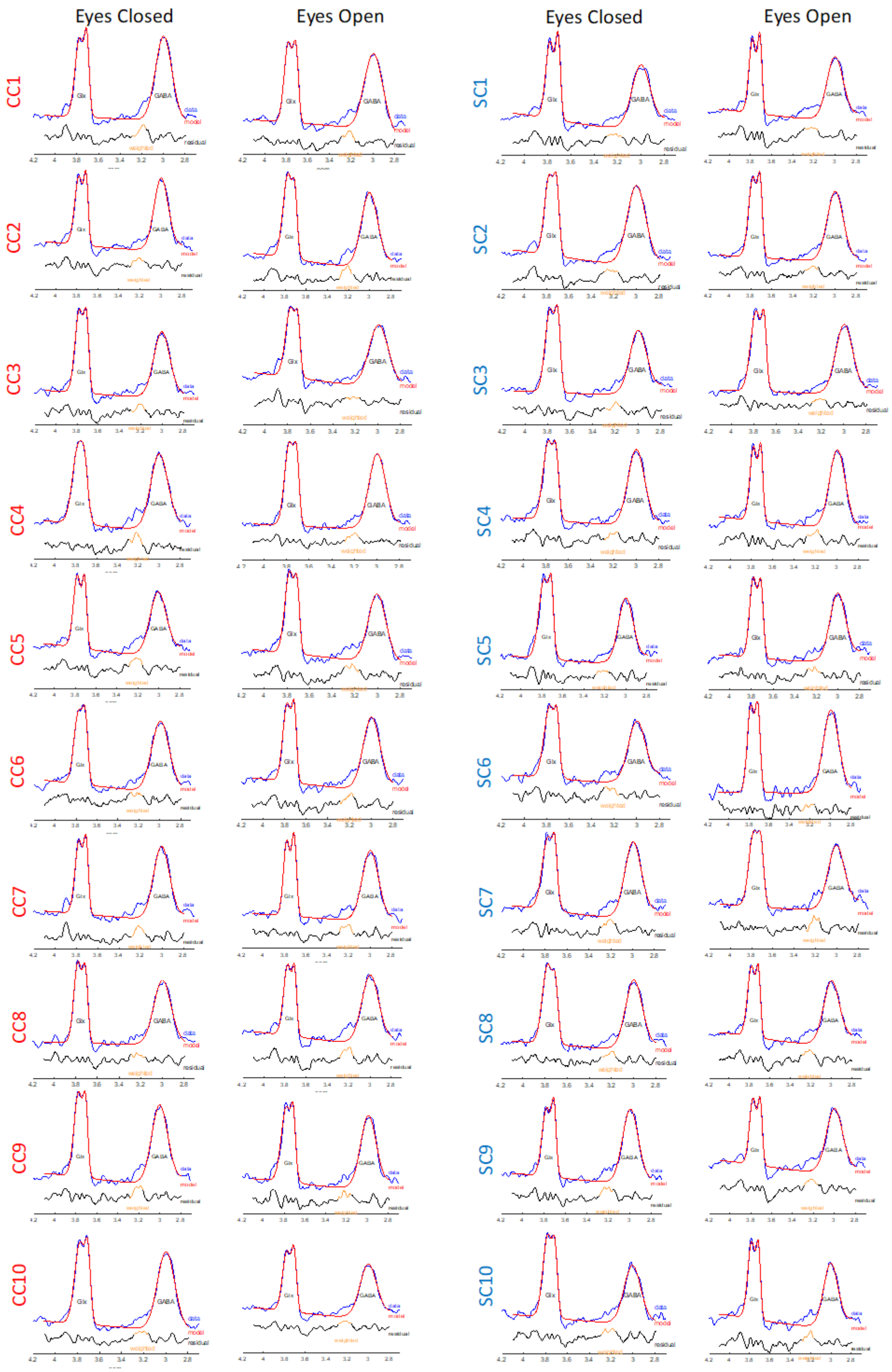


***Figure S8: Edited spectra of participants showing GABA+ and Glx peaks.*** *Individual participants' edited spectra and the respective model fits for congenital cataract-reversal (CC, left) and normally sighted*

*control (SC, right) individuals. Spectra are shown as output by GannetFit.m for the eyes closed and eyes open conditions for each subject.*

**S9. Tests for normality and homogeneity of data**

| **Dependent Variable** | **Shapiro-Wilk Test for Normality** | | | | **Levene's Test for Homogeneity of Variance** | |
|---|---|---|---|---|---|---|
| | **CC (W value)** | **CC (p-value)** | **SC (W value)** | **SC (p-value)** | **F(1,18)** | **p-value** |
| Aperiodic Intercept (EO) | 0.86 | 0.076 | 0.98 | 0.975 | 0.48 | 0.499 |
| Aperiodic Slope (EO) | 0.94 | 0.574 | 0.92 | 0.363 | 0.71 | 0.411 |
| Aperiodic Intercept (EC) | 0.84 | 0.050 | 0.96 | 0.761 | 0.15 | 0.700 |
| Aperiodic Slope (EC) | 0.88 | 0.141 | 0.93 | 0.431 | 0.64 | 0.434 |
| Aperiodic Intercept (LU) | 0.94 | 0.526 | 0.95 | 0.650 | 0.00 | 0.993 |
| Aperiodic Slope (LU) | 0.96 | 0.810 | 0.88 | 0.121 | 0.15 | 0.700 |
| GABA+ (EC) | 0.94 | 0.515 | 0.90 | 0.204 | 2.27 | 0.149 |
| GABA+ (EO) | 0.97 | 0.913 | 0.97 | 0.864 | 0.60 | 0.450 |
| Glx (EC) | 0.97 | 0.881 | 0.87 | 0.102 | 0.69 | 0.416 |
| Glx (EO) | 0.95 | 0.720 | 0.97 | 0.906 | 3.38 | 0.083 |
| GABA+/Glx (EC) | 0.96 | 0.834 | 0.94 | 0.501 | 2.07 | 0.172 |
| GABA+/Glx (EO) | 0.92 | 0.347 | 0.89 | 0.173 | 1.51 | 0.235 |

***Table S9:*** *Results from the Shapiro-Wilk test for normality within each group and Levene's test for homogeneity of variance across groups, for all dependent variables in the reported analyses. The assumption of normality or homogeneity of variance was rejected if p was smaller than 0.05.*

**S10. Rejected epochs from electroencephalography data**

| Group | Eyes Open | Eyes Closed | Visual Stimulation |
|---|---|---|---|
| CC | 13% | 3.1% | 0.2% |
| SC | 18.37% | 2% | 0.1% |

***Table S10:*** *Mean percentage of rejected epochs in each condition for the congenital cataract reversal (CC) and normally sighted control (SC) groups.*

### S11: Individual subjects' aperiodic fits

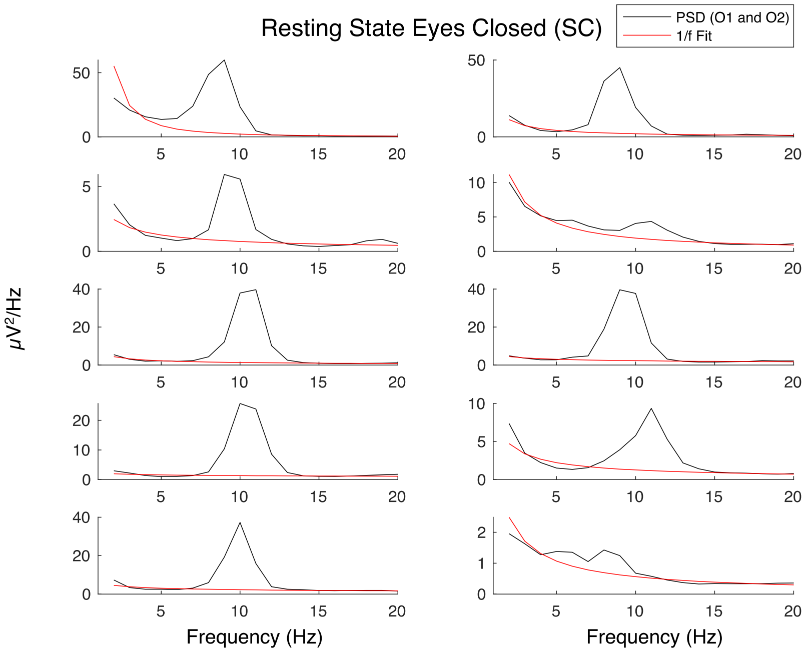


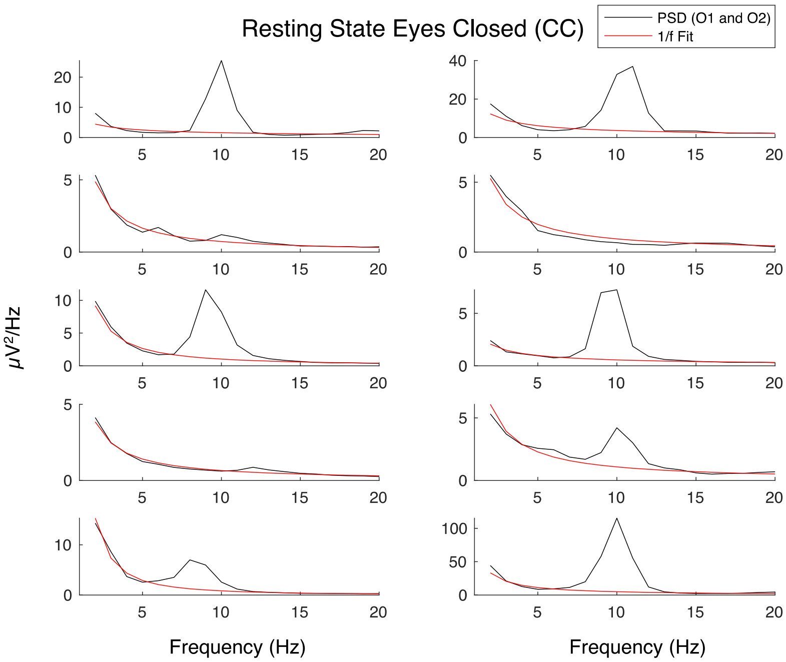


***Figure S11.1: Aperiodic fits for normally sighted control (SC, top) and congenital cataract-reversal (CC, bottom) individuals at occipital electrodes during rest with eyes closed.*** *Solid black lines indicate the power spectral density, red lines indicate the aperiodic (1/f) fit in the 1-20 Hz range, excluding alpha frequencies.*

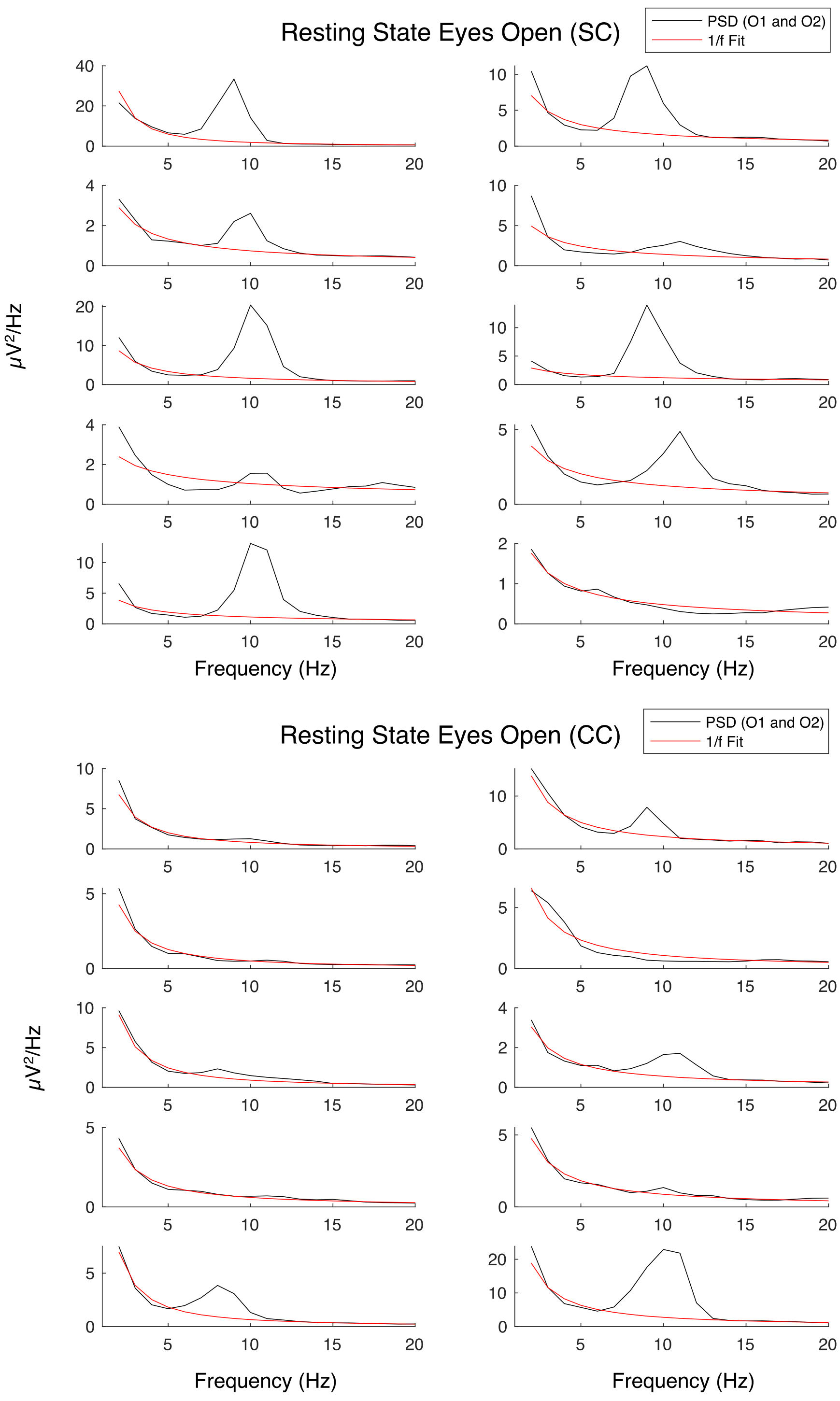


***Figure S11.2: Aperiodic fits for normally sighted control (SC, top) and congenital cataract-reversal (CC, bottom) individuals at occipital electrodes during rest with eyes open.*** *Solid black lines indicate the power spectral density, red lines indicate the aperiodic (1/f) fit in the 1-20 Hz range, excluding alpha frequencies.*

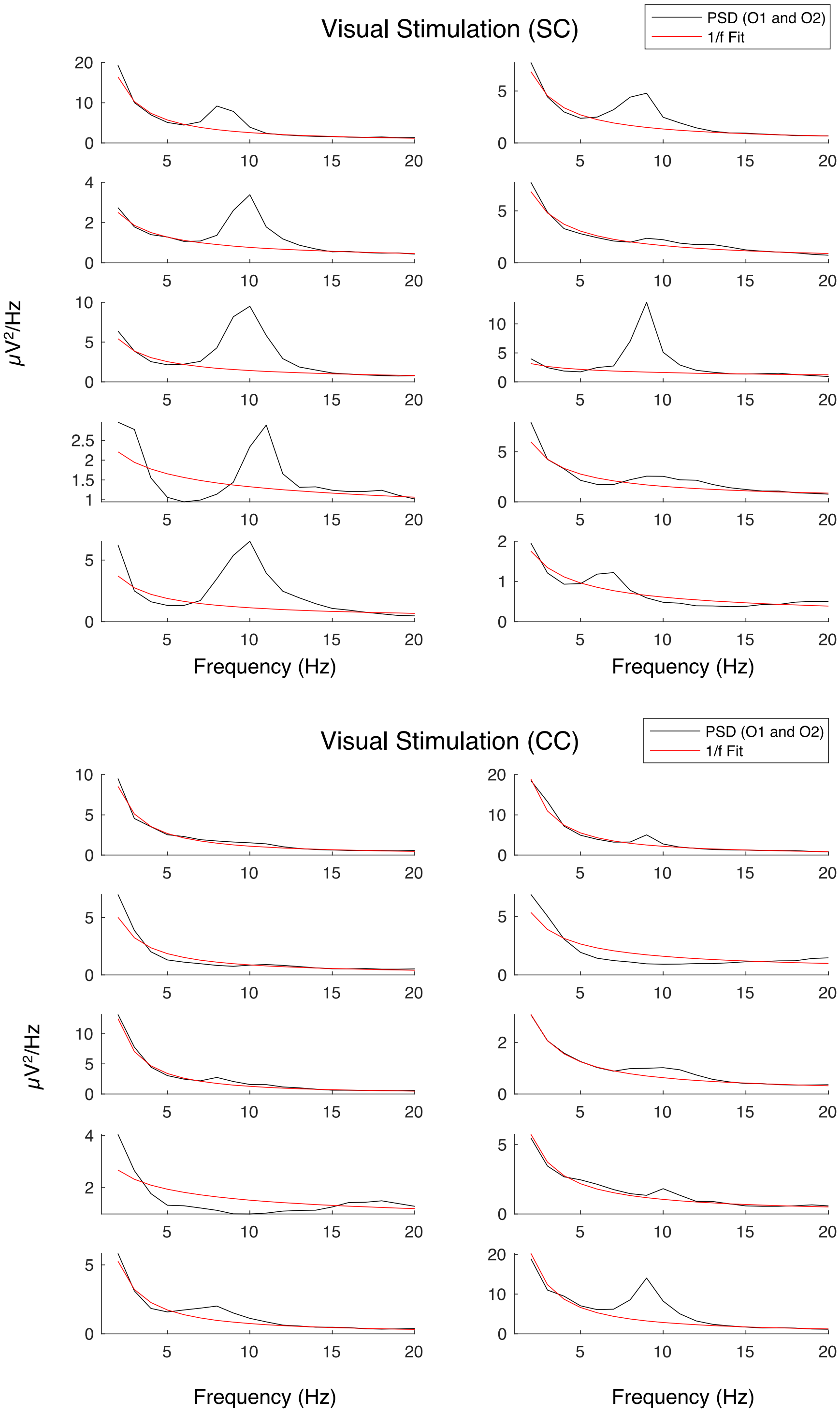


***Figure S11.3: Aperiodic fits for normally sighted control (SC, top) and congenital cataract-reversal (CC, bottom) individuals at occipital electrodes during visual stimulation.** Solid black lines indicate the power spectral density, red lines indicate the aperiodic (1/f) fit in the 1-20 Hz range, excluding alpha frequencies.*

| | Eyes Closed (EC) | Eyes Open (EO) | Visual Stimulation (LU) |
|---|---|---|---|
| Sighted Control (SC) | 0.96 | 0.96 | 0.91 |
| Congenital Cataract Reversal (CC) | 0.95 | 0.98 | 0.99 |

***Table S11:*** *Goodness of fit ($R^2$) values for the EEG aperiodic spectrum. Average $R^2$ values are reported for each group and condition.*

### S12. Exploratory correlation analysis between Magnetic Resonance Spectroscopy measures and visual deprivation history

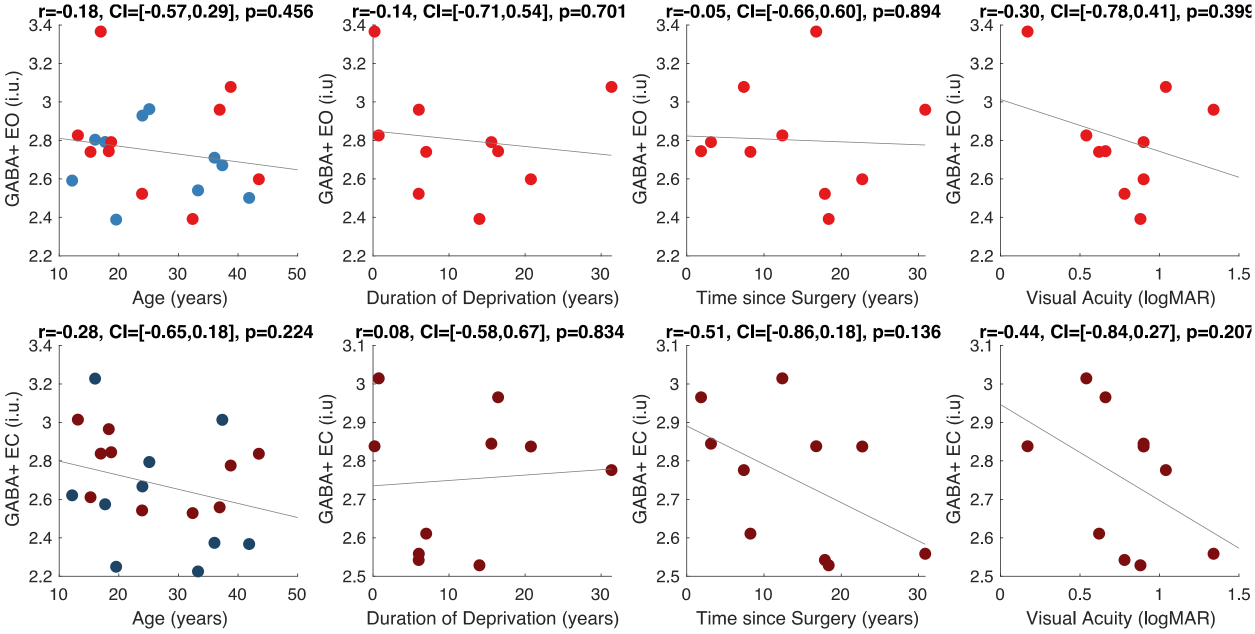


***Figure S12.1. Effect of visual deprivation history on GABA+ concentration:*** *Correlations between visual cortex GABA+ concentration and chronological age of the congenital cataract-reversal (CC, red) and normally sighted individuals (SC, blue, see left panel). Second to fourth panels depict correlations between visual cortex GABA+ concentration and duration of visual deprivation, time since surgery and visual acuity in the CC individuals, respectively. Correlations were separately calculated for the eyes open (EO, top row) and eyes closed (EC, bottom row) conditions. The 95% confidence intervals (CI) of the correlation coefficients (r) are reported.*

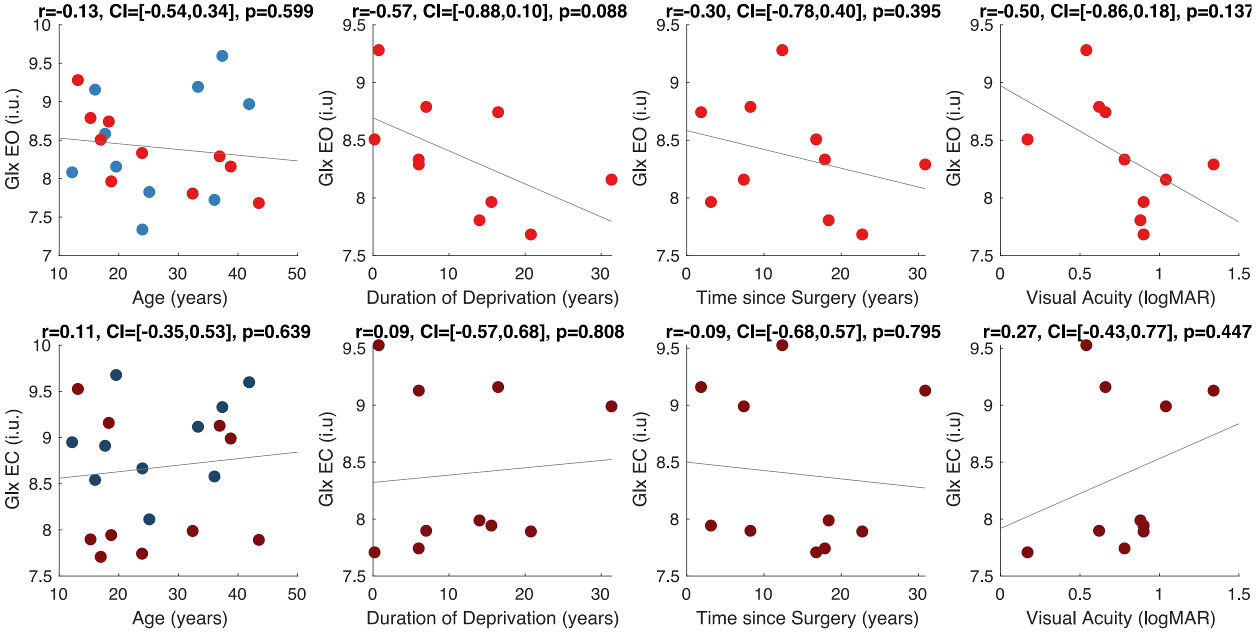


***Figure S12.2. Effect of visual deprivation history on Glx concentration:*** *Correlations between visual cortex Glx concentration and chronological age of the congenital cataract-reversal (CC, red) and normally sighted individuals (SC, blue, see left panel). Second to fourth panels depict correlations between visual cortex Glx concentration and duration of visual deprivation, time since surgery and visual acuity in the CC individuals, respectively. Correlations were separately calculated for the eyes open (EO, top row) and eyes closed (EC, bottom row) conditions. The 95% confidence intervals (CI) of the correlation coefficients (r) are reported.*

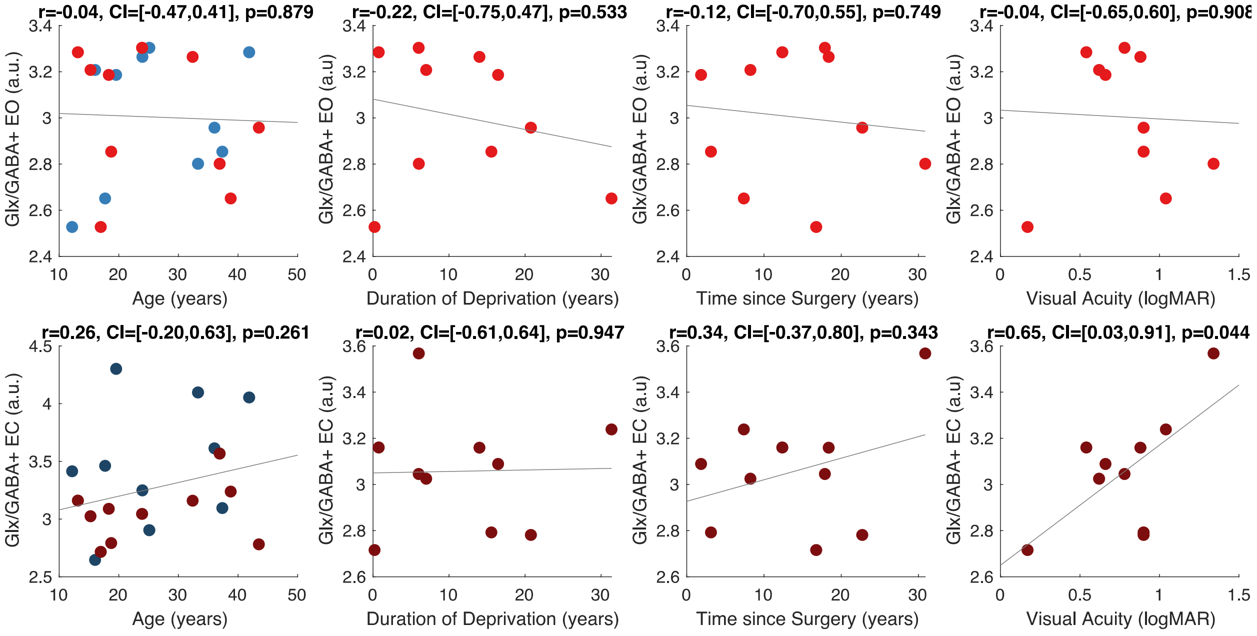

*Figure S12.3. Effect of visual deprivation history on Glx/GABA concentration:* *Correlations between visual cortex Glx/GABA+ concentration and chronological age of the congenital cataract-reversal (CC, red) and normally sighted individuals (SC, blue, see left panel). Second to fourth panels depict correlations between visual cortex Glx/GABA+ concentration and duration of visual deprivation, time since surgery and visual acuity in the CC individuals, respectively. Correlations were separately calculated for the eyes open (EO, top row) and eyes closed (EC, bottom row) conditions. The 95% confidence intervals (CI) of the correlation coefficients (r) are reported.*

**S13. Magnetic Resonance Spectroscopy OFF-Spectra**

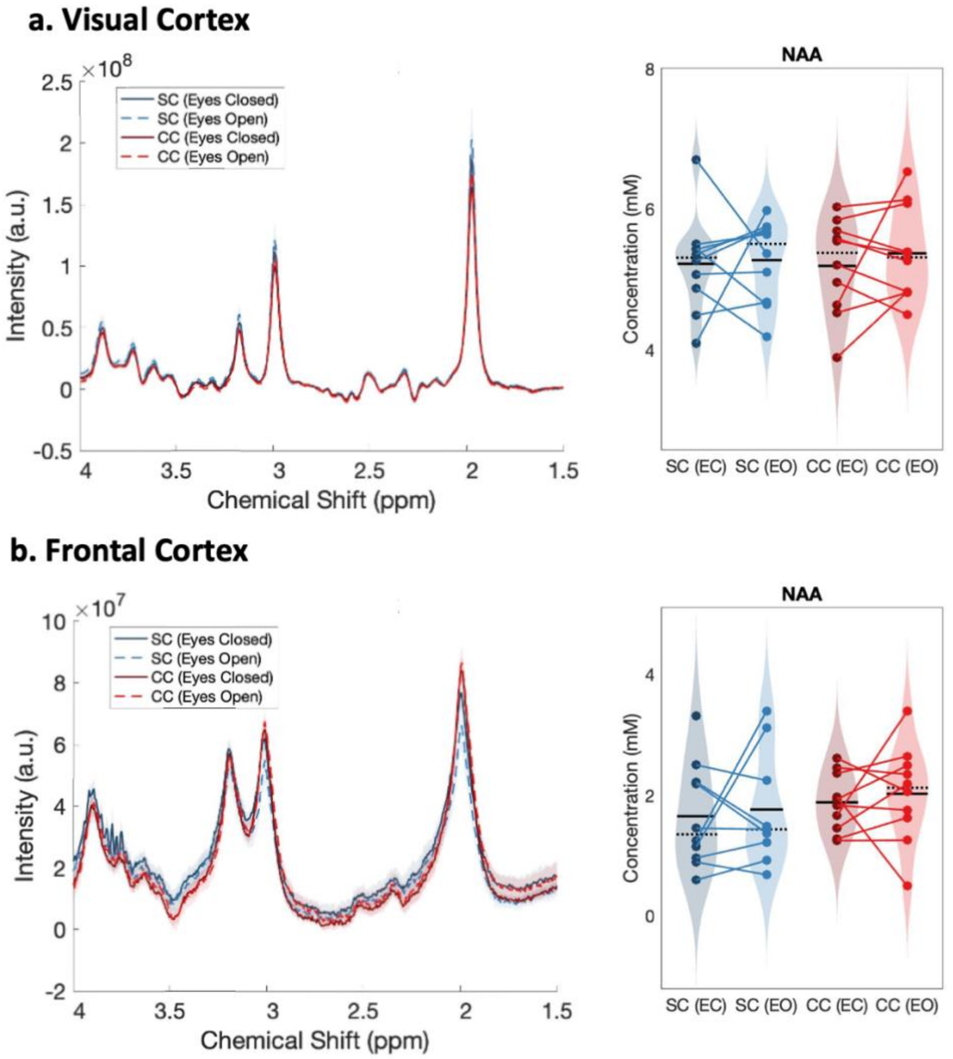


***Figure S13**: **OFF spectra obtained from Magnetic Resonance Spectroscopy (MRS).*** *a. The average spectra show NAA peaks in the visual cortices of normally sighted individuals (SC, green) and individuals*

*with reversed congenital cataracts (CC, red) are shown. Spectra are displayed for the eyes open (EO), and eyes closed (EC) conditions. The standard error of the mean is shaded. NAA concentration distributions for each group and condition are demonstrated as violin plots on the right. The solid black lines indicate mean values, and dotted lines indicate median values. The coloured lines connect values of individual participants across conditions. b. Corresponding average MRS spectra and NAA concentration distributions measured from the frontal cortex are displayed.*

**S14: Aperiodic measures across frontal electrodes**

To assess the spatial specificity of aperiodic EEG measures, we compared the aperiodic slope and intercept calculated across the frontal electrodes FP1 and FP2 between congenital cataract-reversal (CC) and age-matched sighted control individuals (SC). We found that neither group nor condition significantly predicted the aperiodic offset across frontal electrodes (Main effect of group $F(1,59) = 0.11$, $p = 0.746$, Main effect of condition $F(1,59) = 0.14$, $p = 0.712$, Group-by-condition interaction $F(1,59) = 0.05$, $p = 0.885$). Moreover, the aperiodic slope did not vary with either group or condition in frontal electrodes (Main effect of group $F(1,59) = 0.09$, $p = 0.771$, Main effect of condition $F(2,59) = 0.93$, $p = 0.400$, Group-by-condition interaction $F(2,59) = 0.22$, $p = 0.801$) (Figure S14.1).

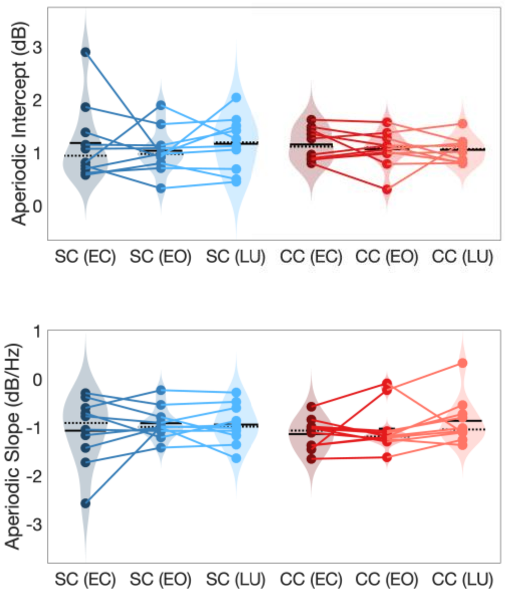


***Figure S14.1: Aperiodic intercept (top) and slope (bottom) for congenital cataract-reversal (CC, red) and age-matched normally sighted control (SC, blue) individuals in frontal electrodes.*** *Distributions of these parameters are displayed as violin plots for three conditions; at rest with eyes closed (EC), at rest*

*with eyes open (EO) and during visual stimulation (LU). Aperiodic parameters were calculated across electrodes Fp1 and Fp2. Solid black lines indicate mean values, dotted black lines indicate median values. Coloured lines connect values of individual participants across conditions.*

### S15. Exploratory correlation analysis between the aperiodic slope and intercept of the EEG power spectrum and visual deprivation history

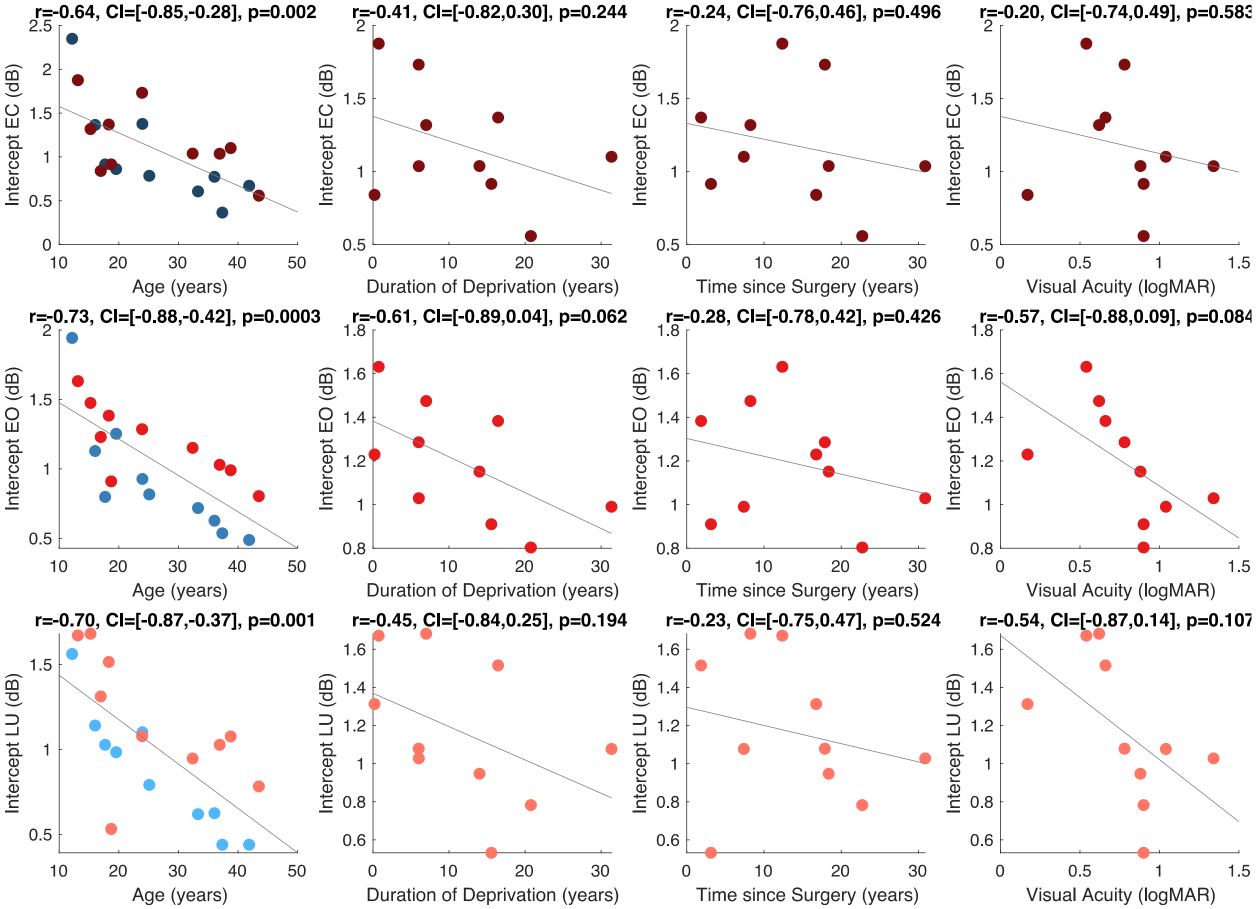


***Figure S15.1: Effect of visual deprivation history on aperiodic intercept:*** *Correlations between aperiodic intercept at occipital electrodes and chronological age of the congenital cataract-reversal (CC, red) and normally sighted individuals (SC, blue, see left panel). Second to fourth panels depict correlations between aperiodic intercept and duration of visual deprivation, time since surgery and visual acuity in the CC individuals, respectively. Correlations were separately calculated for the aperiodic intercept while participants viewed stimuli that changed in luminance (LU, top row) and the eyes open (EO, middle row) and eyes closed (EC, bottom row) conditions. The 95% confidence intervals (CI) of the correlation coefficients (r) are reported.*

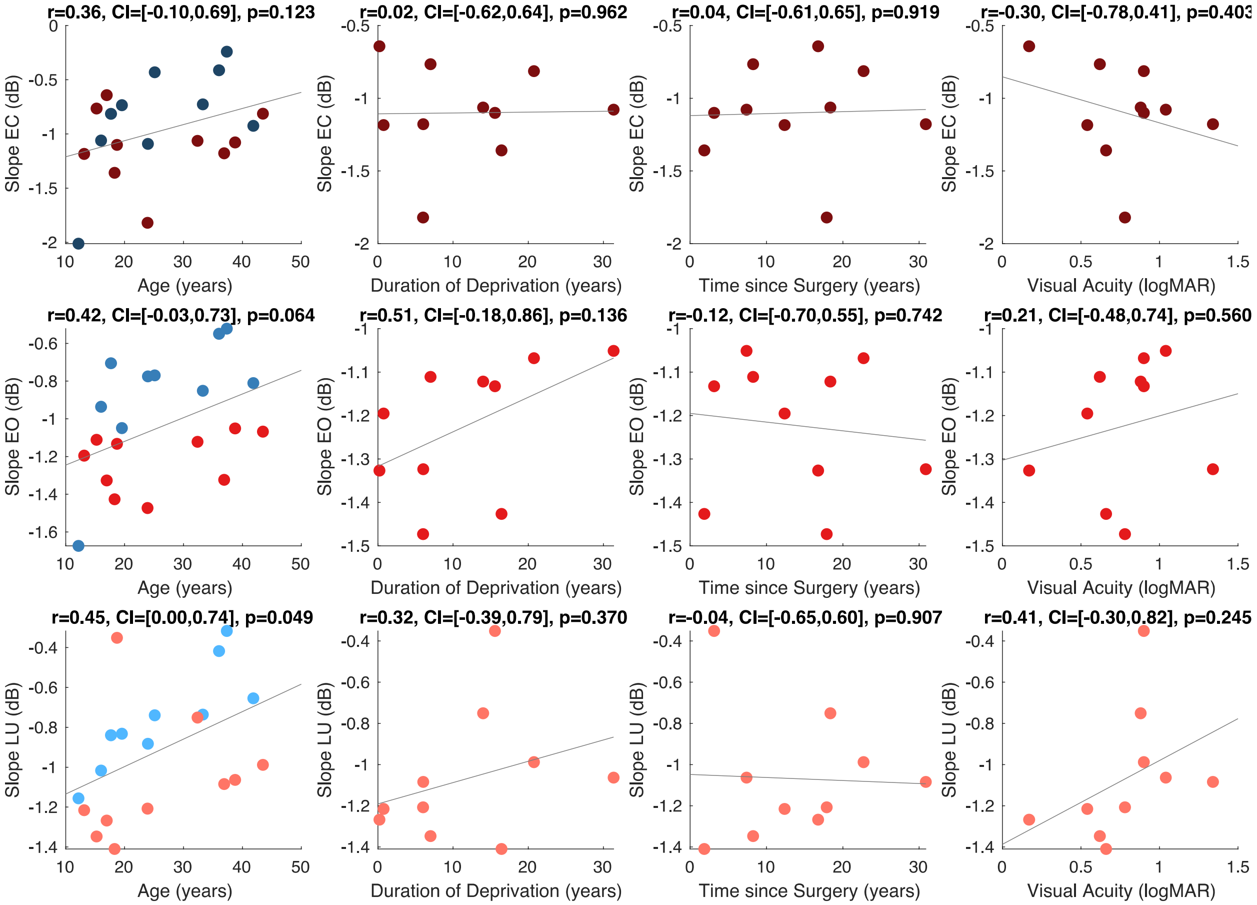


***Figure S15.2: Effect of visual deprivation history on aperiodic slope:*** *Correlations between aperiodic slope across occipital electrodes and chronological age of the congenital cataract-reversal (CC, red) and normally sighted individuals (SC, blue, see left panel). Second to fourth panels depict correlations between aperiodic slope and duration of visual deprivation, time since surgery and visual acuity in the CC individuals, respectively. Correlations separately calculated for the aperiodic slope while participants viewed flickering stimuli (LU, top row) and the eyes open (EO, middle row) and eyes closed (EC, bottom row) conditions. The 95% confidence intervals (CI) of the correlation coefficients (r) are reported.*

**S16. Linear Regression between Glx and aperiodic intercept with age as covariate**

A linear regression was conducted within the CC group to predict the aperiodic intercept during visual stimulation, based on age and visual cortex Glx concentration. The results of the regression analysis indicated that the model explained a significant proportion of the variance in the aperiodic intercept, $R^2$=0.82, $t$(2,7)=16.1, $p$=0.0024. Note that the coefficient for age was not significant, $\beta$=0.007, t(7)=0.82, $p$=0.439. The regression coefficients and their respective statistics are presented in Tables S16.1.

| Predictor | Estimate | SE | t | p |
|---|---|---|---|---|
| Model Intercept | -5.75 | 1.71 | -3.36 | 0.012 |
| Age | 0.007 | 0.008 | 0.82 | 0.439 |
| Glx | 0.81 | 0.19 | 4.36 | 0.003 |

***Table S16.1:*** *Regression summary for the effects of Glutamate/Glutamine (Glx) and age on aperiodic intercept (Visual Stimulation) in the congenital cataract reversal (CC) group in the visual stimulation (LU) condition.*

A second regression was conducted to predict the aperiodic intercept in the CC group during eye opening at rest, based on age and visual cortex Glx concentration. The results of the regression analysis indicated that the model explained a significant proportion of the variance in the aperiodic intercept, $R^2$=0.842, $t$(2,7)=18.6, $p$=0.00159. Note that the coefficient for age was not significant, $\beta$=−0.005, t(7)=−0.90, $p$=0.400. The regression coefficients and their respective statistics are presented in Table S16.2.

| Predictor | Estimate | SE | t | p |
|---|---|---|---|---|
| Model Intercept | -2.07 | 1.11 | -1.86 | 0.106 |
| Age | -0.005 | 0.005 | -0.90 | 0.400 |
| Glx | 0.40 | 0.12 | 3.35 | 0.012 |

***Table S16.2:*** *Regression summary for the effects of Glutamate/glutamine (Glx) concentration and age on aperiodic intercept during eye opening at rest (EO) in the congenital cataract reversal (CC) group*

Given that the Glx coefficient was significant in both models, and age did not significantly predict either outcome. we concluded that Glx predicted the intercept of the aperiodic intercept.

**S17. Exploratory correlation analysis between Electroencephalography and Magnetic Resonance Spectroscopy measures**

We tested the correlations between Glx, GABA+ and Glx/GABA+ measured at rest, and EEG aperiodic broadband intercept as well as slope in CC and SC individuals, measured at rest and while participants observed a flickering visual stimulus. Below, we report the exploratory correlations prior to Bonferroni correction for 6 comparisons (Figures S17.1 – S17.5). Note that the correlation found between the aperiodic slope (1-20 Hz) and Glx concentration (see Figure S17.5) was not significant (all p's > 0.219) after correcting for multiple comparisons.

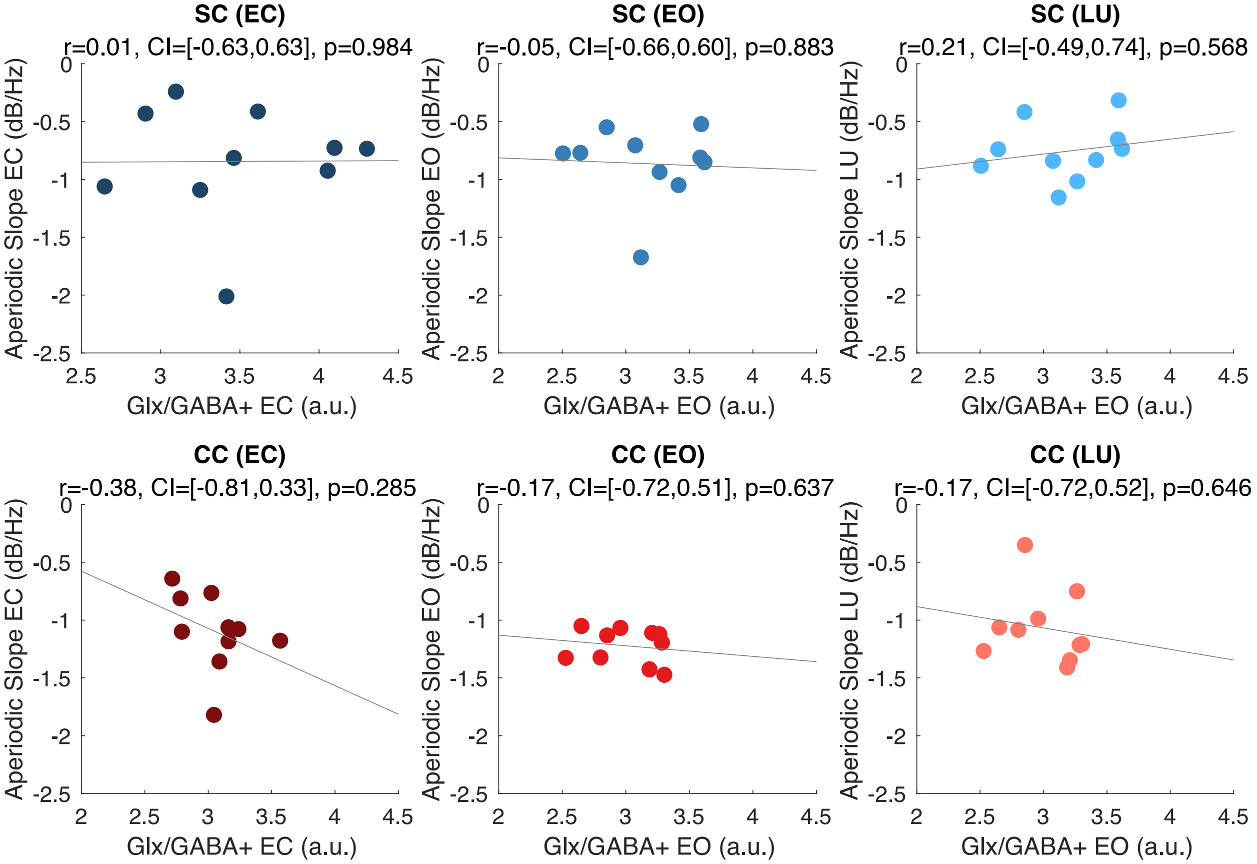


***Figure S17.1: Correlation between aperiodic slope and Glx/GABA+ concentration.*** *Correlations between the aperiodic slope and visual cortex Glx/GABA+ concentration measured at rest with eyes closed (EC) (left panels) and eyes open (EO) (middle panels), and the correlation between aperiodic slope measured while subjects viewed flickering stimuli (LU) and visual cortex Glx/GABA+ concentration measured in the EO condition (right panels), are depicted. Correlations were calculated separately for normally sighted control (SC, blue, top row) and congenital cataract-reversal (CC, red, bottom row) individuals. The 95% confidence intervals (CI) of the correlation coefficients (r) are reported.*

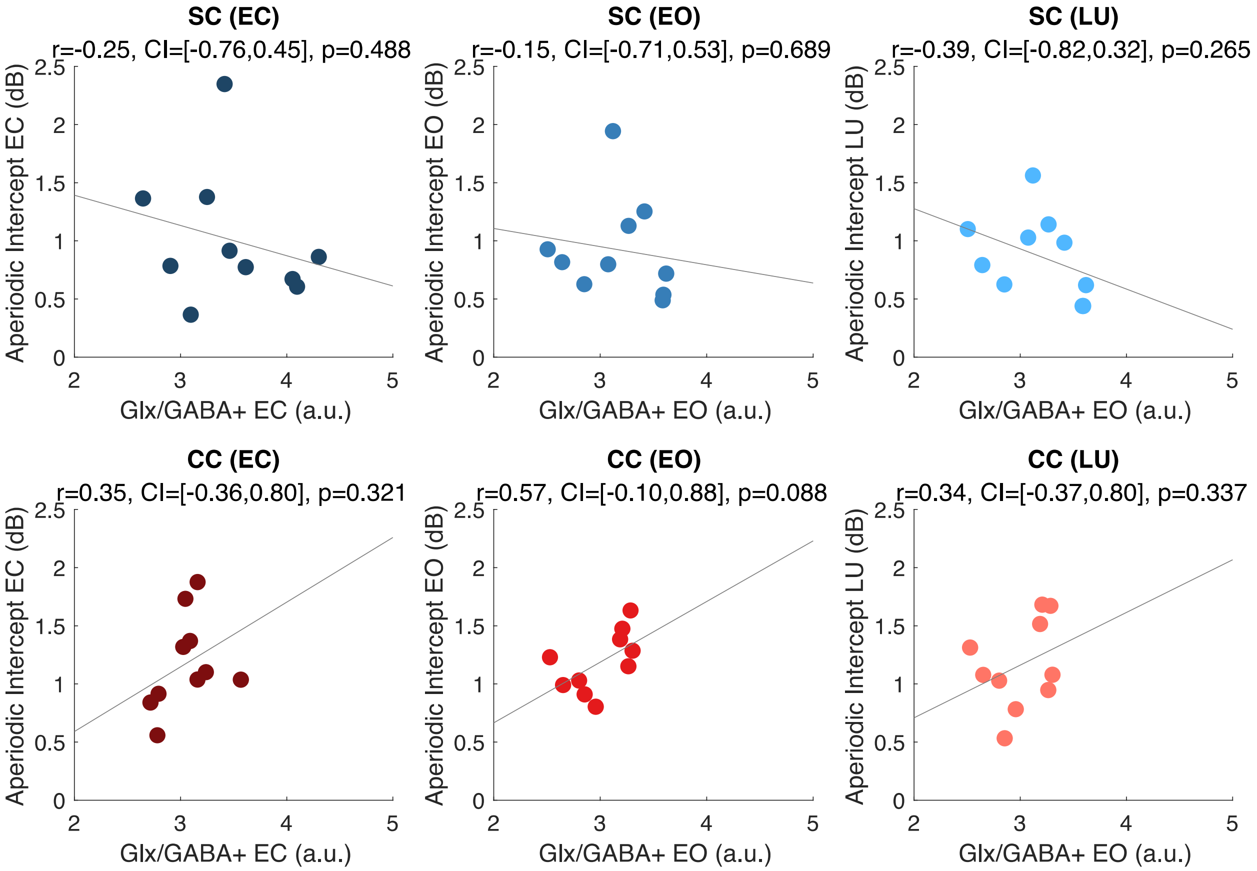


***Figure S17.2: Correlation between aperiodic intercept and Glx/GABA+ concentration.*** *Correlations between the aperiodic intercept and visual cortex Glx/GABA+ concentration measured at rest with eyes closed (EC) (left panels) and eyes open (EO) (middle panels), and the correlation between aperiodic intercept measured while subjects viewed flickering stimuli (LU) and visual cortex Glx/GABA+ concentration measured in the EO condition (right panels), are depicted. Correlations were calculated separately for normally sighted control (SC, blue, top row) and congenital cataract-reversal (CC, red, bottom row) individuals. The 95% confidence intervals (CI) of the correlation coefficients (r) are reported.*

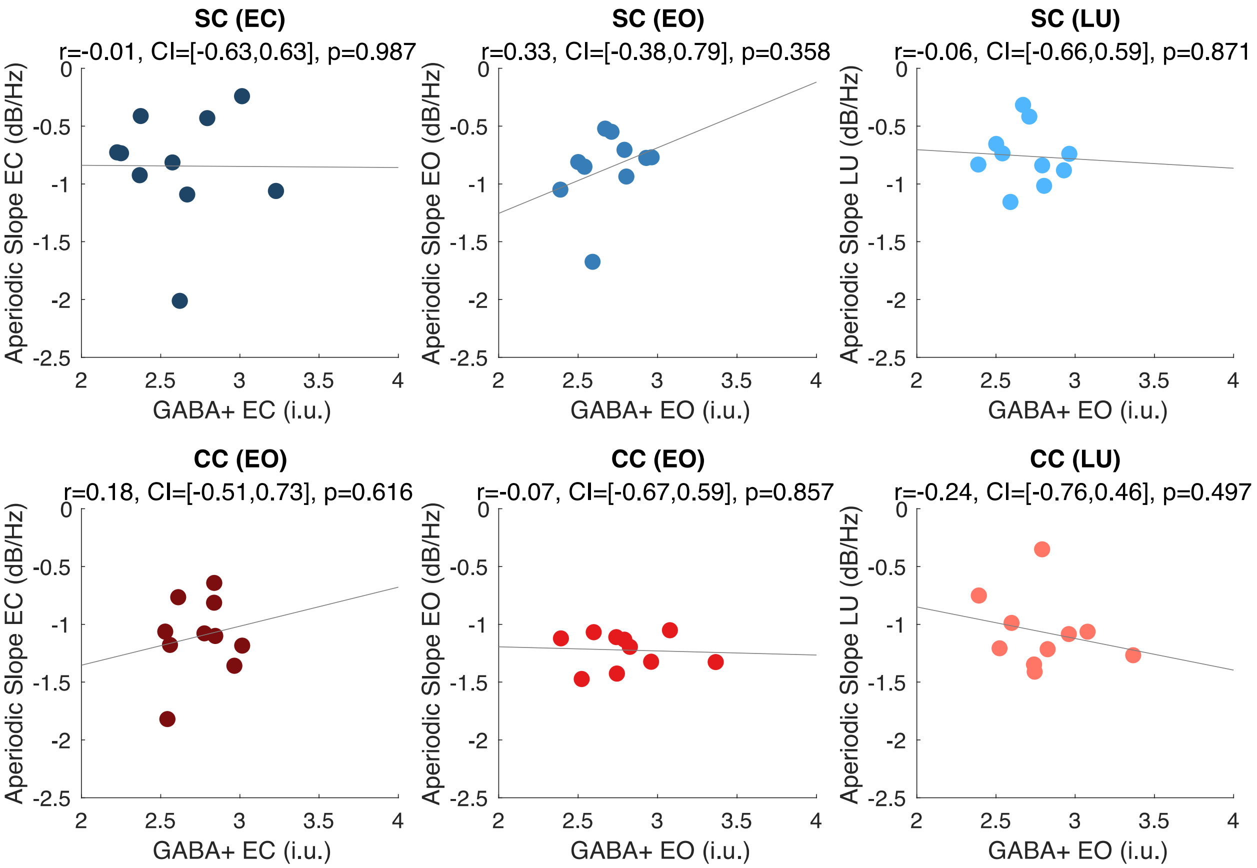


***Figure S17.3: Correlation between aperiodic slope and GABA+ concentration.*** *Correlations between the aperiodic slope and visual cortex GABA+ concentration measured at rest with eyes closed (EC) (left panels) and eyes open (EO) (middle panels), and the correlation between aperiodic slope measured while subjects viewed flickering stimuli (LU) and visual cortex GABA+ concentration measured in the EO condition (right panels), are depicted. Correlations were calculated separately for normally sighted control (SC, blue, top row) and congenital cataract-reversal (CC, red, bottom row) individuals. The 95% confidence intervals (CI) of the correlation coefficients (r) are reported.*

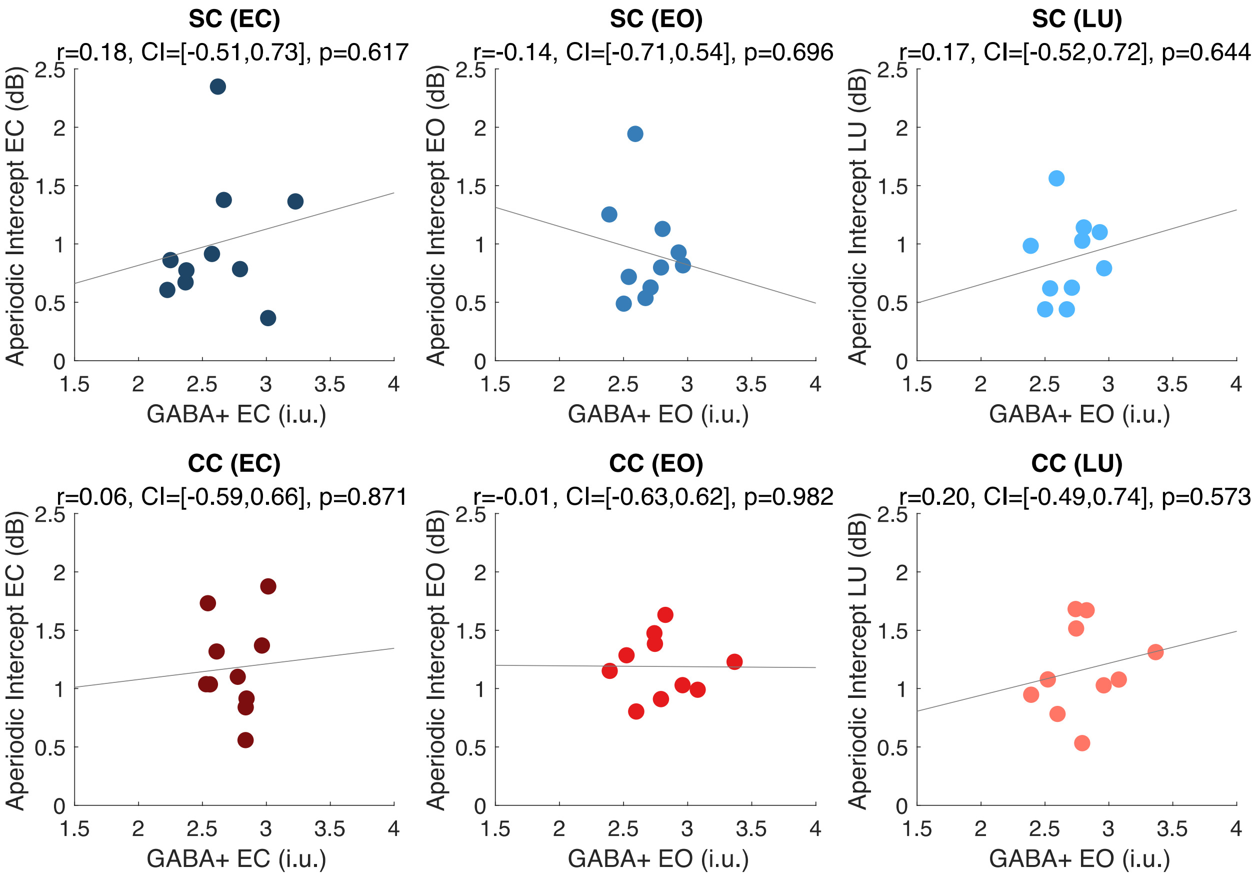


***Figure S17.4: Correlation between aperiodic intercept and GABA+ concentration.*** *Correlations between the aperiodic intercept and visual cortex GABA+ concentration measured at rest with eyes closed (EC) (left panels) and eyes open (EO) (middle panels), and the correlation between aperiodic intercept measured while subjects viewed flickering stimuli (LU) and visual cortex GABA+ concentration measured in the EO condition (right panels), are depicted. Correlations were calculated separately for normally sighted control (SC, blue, top row) and congenital cataract-reversal (CC, red, bottom row) individuals. The 95% confidence intervals (CI) of the correlation coefficients (r) are reported.*

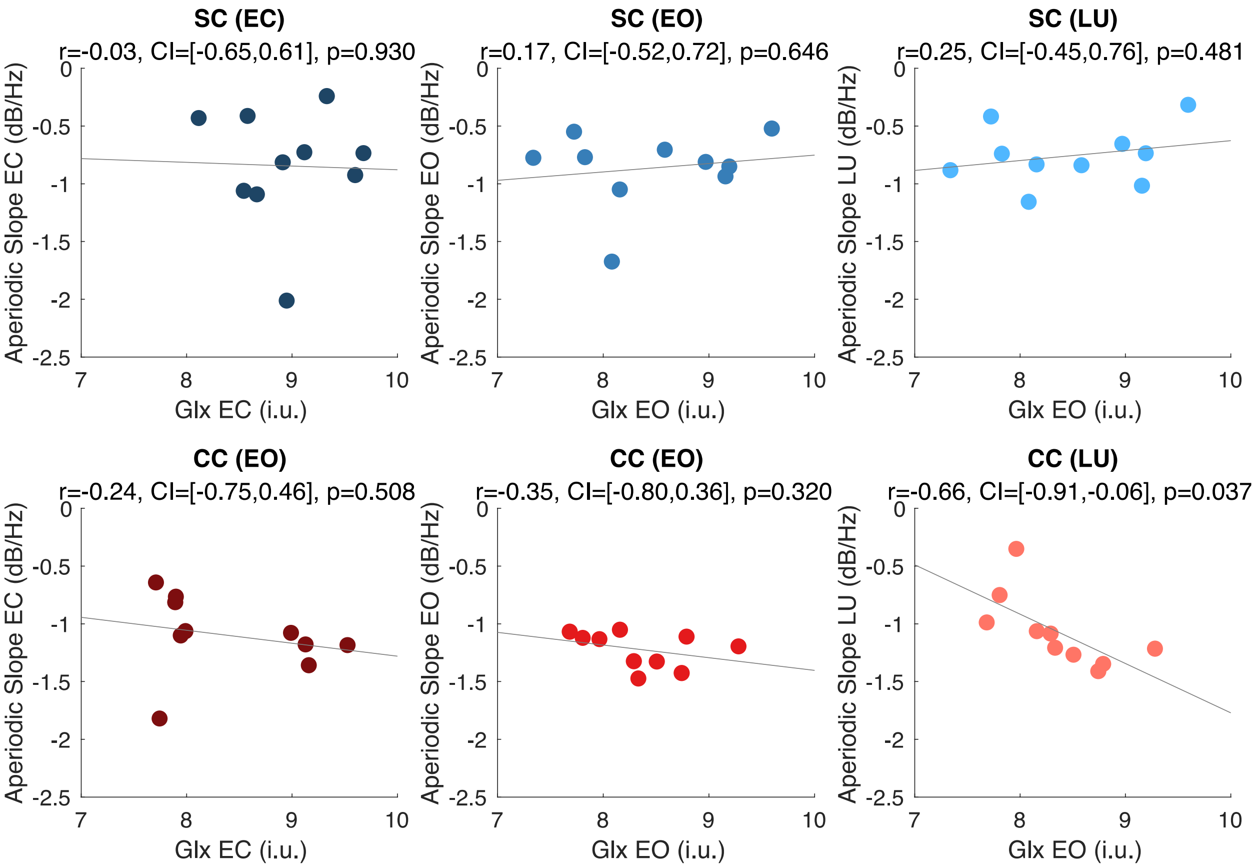


***Figure S17.5: Correlation between aperiodic slope and Glx concentration.*** *Correlations between the aperiodic slope and visual cortex Glx concentration measured at rest with eyes closed (EC) (left panels) and eyes open (EO) (middle panels), and the correlation between aperiodic slope measured while subjects viewed flickering stimuli (LU) and visual cortex Glx concentration measured in the EO condition (right panels), are depicted. Correlations were calculated separately for normally sighted control (SC, blue, top row) and congenital cataract-reversal (CC, red, bottom row) individuals. The 95% confidence intervals (CI) of the correlation coefficients (r) are reported.*

**S18: Correspondence of 1-20 Hz findings with Ossandón et al., 2023**

The resting-state EEG data from the 10 congenital cataract reversal (CC) individuals in the present study corresponded to that of 28 additional CC subjects tested by Ossandón et al., 2023.

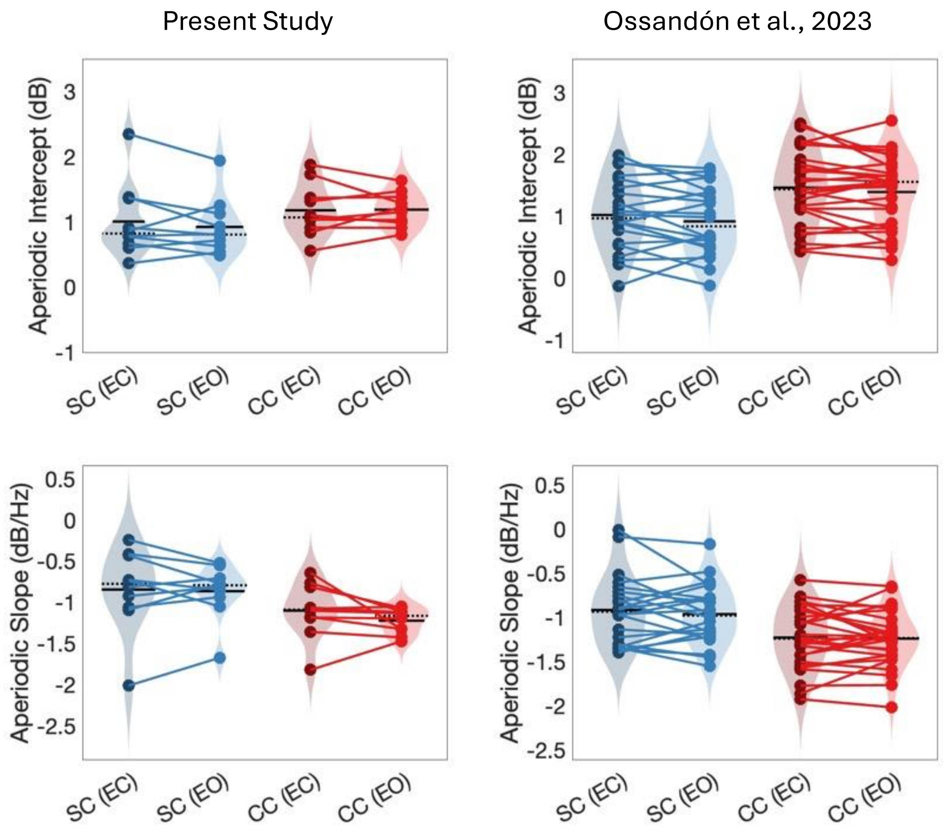


***Figure S18: Aperiodic offset and slope in the 1-20 Hz range from occipital electrodes in congenital cataract reversal (CC) and normally sighted control (SC) individuals of the present study (left) and additional 28 subjects of Ossandon et al. (2023).*** *Aperiodic intercepts (Top) and slope (bottom) distributions for each group and condition are displayed as violin plots. Solid black lines indicate mean values, dotted black lines indicate median values. Coloured lines connect values of individual participants across conditions.*

**S19. Alpha amplitude compared between congenital cataract-reversal and sighted control individuals**

This dataset is a subset of prior findings of reduced alpha amplitude in congenital cataract-reversal (CC) vs normally sighted control individuals (SC) (Ossandón et al., 2023; Pant et al., 2023). We tested for differences in alpha amplitude between the 10 CC individuals of the MRS study and their controls and replicated the results of Ossandon et al. and Pant et al., in the present sample (Figure S19). An ANOVA revealed that the alpha amplitude was lower in CC than in SC individuals across conditions (main effect of group: $F(1,59) = 8.95$, $p = 0.004$, $\eta_p^2 = 0.14$, group-by-condition interaction: $F(2,59) = 0.8$, $p = 0.454$, $\eta_p^2 = 0.03$). As expected, eye closure increased alpha activity compared to eye opening and visual stimulation (main effect of condition: $F(2,59) = 13.12$, $p < 0.001$, $\eta_p^2 = 0.33$).

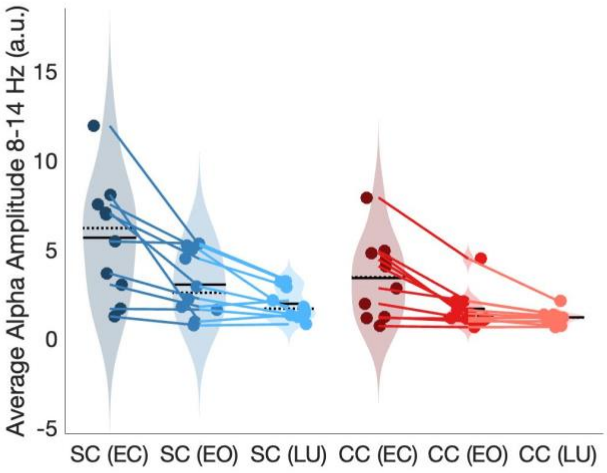


*Figure S19:* ***Aperiodic-corrected alpha amplitude in congenital cataract-reversal and normally sighted individuals.*** *Aperiodic-corrected alpha amplitudes (8-14 Hz) distributions for each group and condition are displayed as violin plots. Solid black lines indicate mean values, dotted black lines indicate median values. Coloured lines connect values of individual participants across conditions.*
